# FABP4 as a Therapeutic Host Target Controlling SARS-CoV2 Infection

**DOI:** 10.1101/2024.02.10.579717

**Authors:** Hatoon Baazim, Emre Koyuncu, Gürol Tuncman, M. Furkan Burak, Lea Merkel, Nadine Bahour, Ezgi Simay Karabulut, Grace Yankun Lee, Alireza Hanifehnezhad, Zehra Firat Karagoz, Katalin Földes, Ilayda Engin, Ayse Gokce Erman, Sidika Oztop, Nazlican Filazi, Buket Gul, Ahmet Ceylan, Ozge Ozgenc Cinar, Fusun Can, Hahn Kim, Ali Al-Hakeem, Hui Li, Fatih Semerci, Xihong Lin, Erkan Yilmaz, Onder Ergonul, Aykut Ozkul, Gökhan S. Hotamisligil

## Abstract

Host metabolic fitness is a critical determinant of infectious disease outcomes. Obesity, aging, and other related metabolic disorders are recognized as high-risk disease modifiers for respiratory infections, including coronavirus infections, though the underlying mechanisms remain unknown. Our study highlights fatty acid-binding protein 4 (FABP4), a key regulator of metabolic dysfunction and inflammation, as a modulator of SARS-CoV-2 pathogenesis, correlating strongly with disease severity in COVID-19 patients. We demonstrate that loss of FABP4 function, by genetic or pharmacological means, reduces SARS-CoV2 replication and disrupts the formation of viral replication organelles in adipocytes and airway epithelial cells. Importantly, FABP4 inhibitor treatment of infected hamsters diminished lung viral titers, alleviated lung damage and reduced collagen deposition. These findings highlight the therapeutic potential of targeting host metabolism in limiting coronavirus replication and mitigating the pathogenesis of infection.

## Introduction

Viral infections are highly disruptive events that mobilize extensive resources to facilitate replication, organelle remodeling ^1,2^ and immune responses ^3^. The resulting immunometabolic interactions substantially modify host metabolism, while also being greatly influenced by it ^4^. The importance of the host’s metabolic state in determining infectious disease outcomes has been well established, particularly in conditions such as obesity, diabetes, and aging, which substantially increase morbidity and mortality during coronavirus infections even in vaccinated individuals ^5,6^. However, a mechanistic understanding of how metabolism affects viral replication and pathogenesis remains limited. In the context of obesity, current hypotheses attribute this heightened pathogenesis to the underlying metabolic dysregulation and low-grade inflammation ^7,8^, which could impair anti-viral immune responses and cause airway hyperresponsiveness ^9^. It is also possible that ectopic lipid deposition and excess nutrient flux, resulting from uncontrolled adipose tissue lipolysis and insulin resistance ^10^, could create an environment more conducive to viral replication. Although studies implementing mouse-adapted viral strains of the Middle East respiratory syndrome coronavirus (MERS-CoV) and SARS-CoV2 reported increased mortality despite comparable lung viral titers across lean and obese high fat diet-fed mice ^11,12^.

FABP4 is an exceptionally versatile regulator of energy resources ^13,14^ and a modulator of metabolic and inflammatory responses ^15,16^. Our current understanding of FABP4 biology attributes its versatility to its dual function, as an intracellular lipid binding chaperon, delivering a wide range of ligands to phospholipid membranes or lipid-sensing receptors such as PPARγ ^17,18^, as well as a secreted hormone capable of interacting with other proteins, such as NDPK and ADK^13^. While there is no known direct link between FABP4 and coronavirus infections, FABP4 is known to promote inflammation and metabolic dysfunction in several comorbidities that constitute a high-risk for coronavirus infection, including diabetes ^13^, cardiovascular disease ^19^, and airway disease ^15,16^. Additionally, its regulation of *de-novo* lipogenesis ^20^, lipid composition ^21^ and intracellular lipid trafficking makes it a promising candidate in facilitating SARS-CoV2 induced organelle remodeling and lipid droplet (LD) utilization^22^. Adipose tissue dysfunction is another common link amongst COVID-19 high-risk comorbidities through its regulation of systemic metabolism and its ability to secrete proinflammatory cytokines ^7^. Adipocytes are permissible to SARS-CoV2 infection ^23,24^, and the detection of viral RNA in the adipose tissue of deceased COVID-19 patients and infected animals ^23–26^ indicates that viral dissemination into the adipose tissue may occur naturally. Adipocytes are the most abundant source of FABP4 ^27^, and together with macrophages, endothelial and epithelial cells, present an overlapping target for SARS-CoV2 infection and FABP4 action. These observations prompted us to directly assess the impact of FABP4 in COVID-19.

In this study, we observed significantly elevated FABP4 levels in the serum and lungs of COVID-19 patients that highly correlate with disease severity. We then examined the intracellular interaction between FABP4 and SARS-CoV2 using cultured adipocytes, and bronchial epithelial cells. We demonstrated that, FABP4 is recruited to the double-membrane vesicles (DMVs) of virus replication organelles (ROs), which are membrane-bound compartments derived from remodeled cellular organelles ^1,2^. Pharmacological inhibition, or genetic deletion of FABP4 resulted in disruption of DMV numbers and organization in SARS-CoV2 infected cells, and broadly impaired coronavirus propagation across various cell types infected with different SARS-CoV2 strains and the common cold coronavirus OC43 (HCoV-OC43). Importantly, FABP4 inhibition in infected hamsters significantly reduced viral titers, and ameliorated lung damage and fibrosis. Together, these data indicate that FABP4 facilitates SARS-CoV2 infection and that therapeutic targeting of FABP4 may be a beneficial treatment strategy for mitigating COVID-19 severity and mortality in humans.

## Results

### FABP4 protein levels correlate with disease severity in COVID-19 patients

To assess FABP4’s involvement in SARS-CoV-2 infection, we conducted immunohistochemical staining for FABP4 in lung biopsies obtained from individuals with COVID-19. This revealed a high FABP4 signal in the endothelial cells lining the small vessels of the lung parenchyma (Figure 1A). We also examined circulating levels of FABP4 in two patient cohorts at distinct stages of disease severity defined according to the WHO severity criteria ^28^ and 45 healthy controls (Tables 1). [Cohort 1 (Tables 2), November 2020 – May 2021, n=283, dominant variants: epsilon and alpha ^29^. Cohort 2 (Tables 3) March-May 2020, n=116, dominant variant: original Wuhan strain] . Multiple serum samples were collected from each patient during hospitalization and marked according to the day post-symptom onset. This analysis revealed a progressive increase in FABP4 levels (Figures 1B, 1C and S1A) that corresponded with an increase in several biomarkers of disease severity, including IL-6, C-reactive protein and circulating leukocytes (Figures 1D and S1B to S1D) and a decrease in lymphocytes (Figure S1E). We further interrogated this association by performing a regression analysis that examines FABP4 concentration over time in each patient, adjusting for age, sex, and BMI. This analysis uncovered a strong correlation between increased circulating FABP4 levels and COVID-19 disease severity (Figure 1E). Consistent with these results, we found that higher FABP4 levels also correlated with the need for non-invasive or mechanical ventilation (Figure 1F). We then examined FABP4 levels in severe and critically ill patients across several high-risk disease modifiers where FABP4 is known to promote inflammation and metabolic dysfunction. By doing so, we found that patients with reported comorbidities and those over 50 years of age had elevated FABP4 levels (Figures 1G,1H, S1F and S1G, Tables 2 and 3). Similar results were also found in patients with a BMI equal to or over 30 and those with cardiometabolic conditions (Figures S1H and S1I, Tables 2 and 3). Taken together, this data points to a strong regulation and potential involvement of FABP4 in the pathophysiology of SARS-CoV2 and suggests that the presence of underlying conditions that are regulated by FABP4 could further influence its involvement.

**Figure 1:**
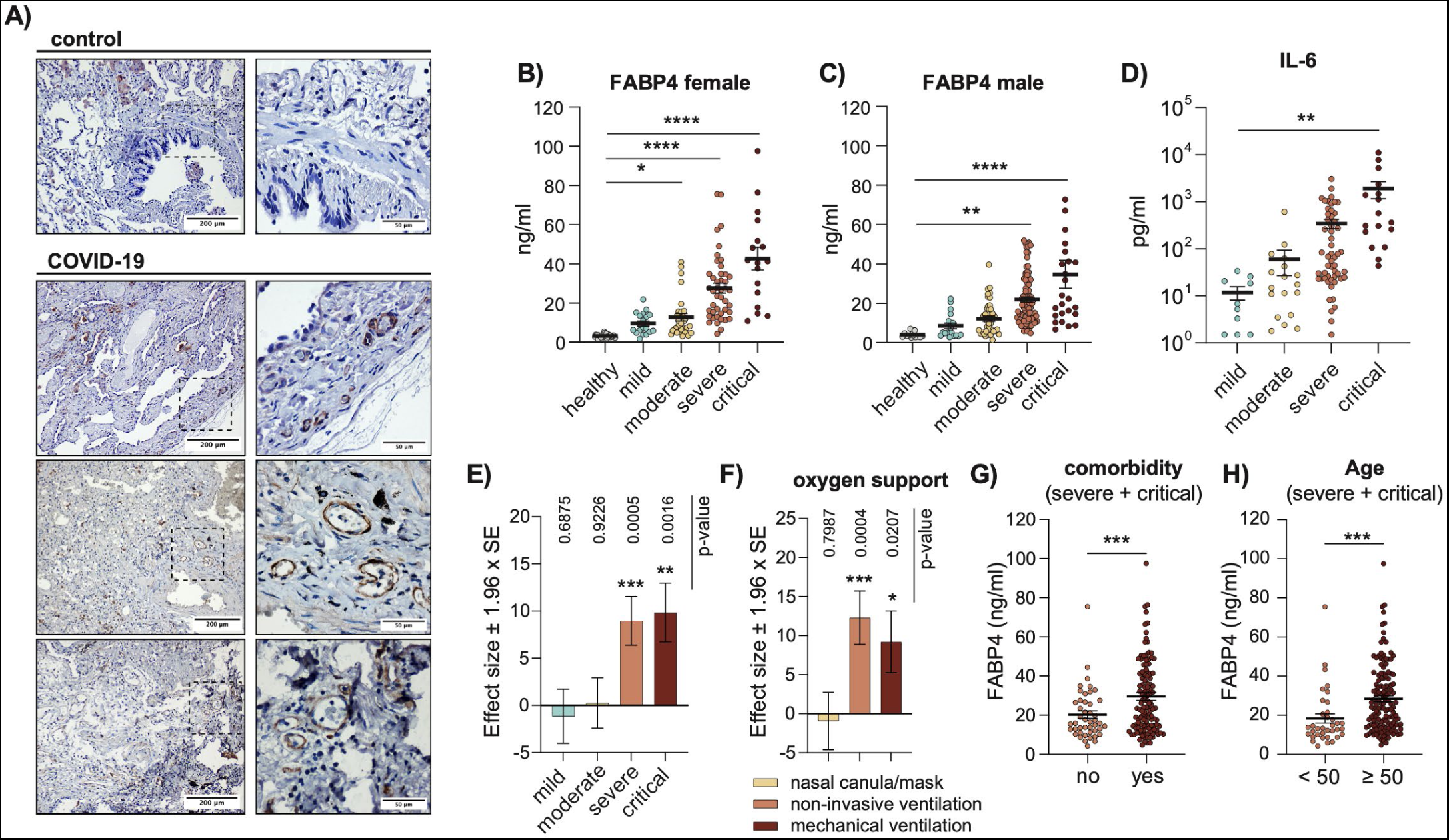
FABP4 levels are increased in the lungs and circulation of COVID-19 patients. A) Immunohistochemical staining of lung biopsies of controls and COVID-19 patients, using anti-FABP4 antibody. B and C) Circulating FABP4 concentrations measured from B) female and C) male COVID-19 patients and healthy controls, stratified based on disease severity. Statistical analysis was performed using one-way ANOVA. D) Circulating IL-6 levels in COVID-19 patients, stratified based on disease severity. Statistical analysis was performed using one-way ANOVA. E and F) Effect size estimates and inference based on regression analysis of FABP4 concentration on E) COVID-19 severity and F) oxygen support measures while accounting for time of collection post symptom onset, age, sex, and BMI, using the linear mixed model to account for the patient-level random effects. E) The healthy controls and, F) patients who did not require oxygen support were used as a reference group. p-values are calculated based on the Wald test. G And H) FABP4 concentration in patients with severe and critical disease, stratified based on G) presence or absence of comorbidities or H) age. Statistical analysis was performed using Welche’s t-test. B to H) Data are derived from patient cohort 1 (collected November 2020 – May 2021). Patients were sampled longitudinally, and the data shown in B, C, D, G and H represent the maximum measured concentration per patient. Data are shown as the mean ± s.e.m.

**Table 1:** Healthy Controls. Cohort 1 (November 2020 – May 2021).

|  | **Total (n=45)** | **Female (n=32)** | **Male (n=13)** |
| --- | --- | --- | --- |
| **Median Age** | 28 (20-56) | 27 (20-56) | 28 (22-41) |
| **Median BMI** | 21.9 (17-30) | 21 (17-28) | 24.6 (21-30) |

**Table 2:** COVID-19 patients. Cohort 1 (November 2020 – May 2021).

|  | **Total (%)** | **COVID-19 Disease severity** |  |  |  |
| --- | --- | --- | --- | --- | --- |
|  |  | **Mild (%)** | **Moderate (%)** | **Severe (%)** | **Critical (%)** |
| **Total Patients** | 283 | 35 (12.4) | 72 (25.4) | 134 (47.3) | 42 (14.8) |
| **Female** | 109 (38.5) | 18 (6.4) | 30 (10.6) | 44 (15.5) | 17 (6) |
| **Male** | 174 (61.5) | 17 (6) | 42 (14.8) | 90 (31.8) | 25 (8.8) |
| **Median Age** | 61 (21– 99) | 53 (21 – 87) | 60 (24 – 83) | 59 (33 – 99) | 74 (54 – 93) |
| **Median BMI** | 27.2 (17.5 – 48.5) | 26 (17.5 – 36.1) | 26.2 (20 – 48.4) | 26.8 (22.6 – 38.5) | 25.9 (24.6 – 39.7) |
| **ICU admission** | 47 (16.6) | 0 (0.0) | 0 (0.0) | 6 (2.1) | 41 (14.5) |
| **Mortality** | 14 (4.9) | 0 (0.0) | 0 (0.0) | 0 (0.0) | 14 (4.9) |
| **All Comorbidity** | 210 (74.2) | 26 (9.2) | 54 (19.1) | 93 (32.9) | 39 (13.8) |
| **Hypertension** | 95 (44.8) | 8 (2.8) | 32 (11.3) | 42 (14.8) | 21 (7.4) |
| **Coronary Artery Disease** | 30 (14.2) | 2 (0.7) | 3 (1.1) | 16 (5.7) | 9 (3.2) |
| **Obesity** | 7 (3.3) | 0 (0.0) | 2 (0.7) | 2 (0.7) | 3 (1.1) |
| **Diabetes** | 57 (26.9) | 6 (2.1) | 14 (4.9) | 29 (10.2) | 8 (2.8) |
| **COPD** | 11 (5.2) | 1 (0.4) | 0 (0.0) | 4 (1.4) | 6 (2.1) |
| **Asthma** | 9 (4.2) | 2 (0.7) | 2 (0.7) | 5 (1.8) | 0 (0.0) |

**Table 3.** COVID-19 patients. Cohort 2 (March – May 2020).

|  | **Total** | **Moderate** | **COVID-19 Disease severity** |  |
| --- | --- | --- | --- | --- |
|  |  |  | **Severe** | **Critical** |
| **Total Patients** | 116 | 52 (44.8) | 21 (18.1) | 42 (36.2) |
| **Female** | 48 (41.4) | 29 (25) | 7 (6) | 12 (10.3) |
| **Male** | 68 (58.6) | 23 (19.8) | 14 (12.1) | 31 (26.7) |
| **Median Age** | 65 (35 – 91) | 63 (18 – 87) | 60 (35 – 77) | 69 (45 – 91) |
| **Median BMI** | 27.4 (19.6 – 55) | 27.6 (18.6 – 55) | 27 (22 – 44) | 28 (19.6 – 52.1) |
| **ICU admission** | 31 (26.7) | 0 (0.0) | 0 (0.0) | 31 (26.7) |
| **Mortality** | 9 (7.8) | 0 (0.0) | 0 (0.0) | 9 (7.8) |
| **All Comorbidity** | 58 (50) | 25 (21.6) | 7 (6) | 26 (22.4) |
| **Hypertension** | 46 (39.7) | 22 (19) | 6 (5.2) | 18 (15.5) |
| **Coronary Artery Disease** | 20 (17.2) | 9 (7.8) | 4 (3.4) | 7 (6) |
| **Obesity** | 8 (6.9) | 1 (0.9) | 1 (0.9) | 6 (5.2) |
| **DM** | 25 (21.6) | 8 (3.4) | 4 (3.4) | 13 (11.2) |
| **COPD** | 1 (0.9) | 0 | 0 | 1 (0.9) |
| **Asthma** | 2 (1.7) | 0 | 0 | 2 (1.7) |

### SARS-CoV2 infection is accelerated in mature adipocytes where FABP4 is recruited to DMVs

Due to the central role of adipocytes in aging and obesity-related metabolic dysfunction, and their high FABP4 abundance ^27^, we investigated the dynamics of SARS-CoV2 infection using two infection titers (MOI: 0.1 or 1, SARS-CoV2 WA1/2020), in the <u>h</u>uman <u>Te</u>lomerase <u>R</u>everse <u>T</u>ranscriptase (hTERT) immortalized preadipocyte cell lines ^30^, before and after differentiation. Interestingly, in both infection titers, viral propagation was significantly higher in mature adipocytes, most notably, viral titers, which exceeded that of preadipocytes by several orders of magnitude (Figures 2A–2C and S2A–S2C). We then examined the infection in mature adipocytes more closely and found that, although viral RNA levels were seemingly unchanged over time, nucleocapsid protein abundance did increase (Figures S2I and S2K). Moreover, infection of mature adipocytes triggered a marked increase in IL-6 secretion, at concentrations proportional to the infection titers (Figures 2D and S2D). Infection of adipocytes at various stages of differentiation revealed a striking gradual increase in viral replication over the course of adipocyte maturation as evident by the increased virus titers and nucleocapsid protein abundance (Figures 2E–2G). As expected, FABP4 abundance also increased during adipocyte differentiation ^31^ (Figures 2F and 2G).

**Figure 2:**
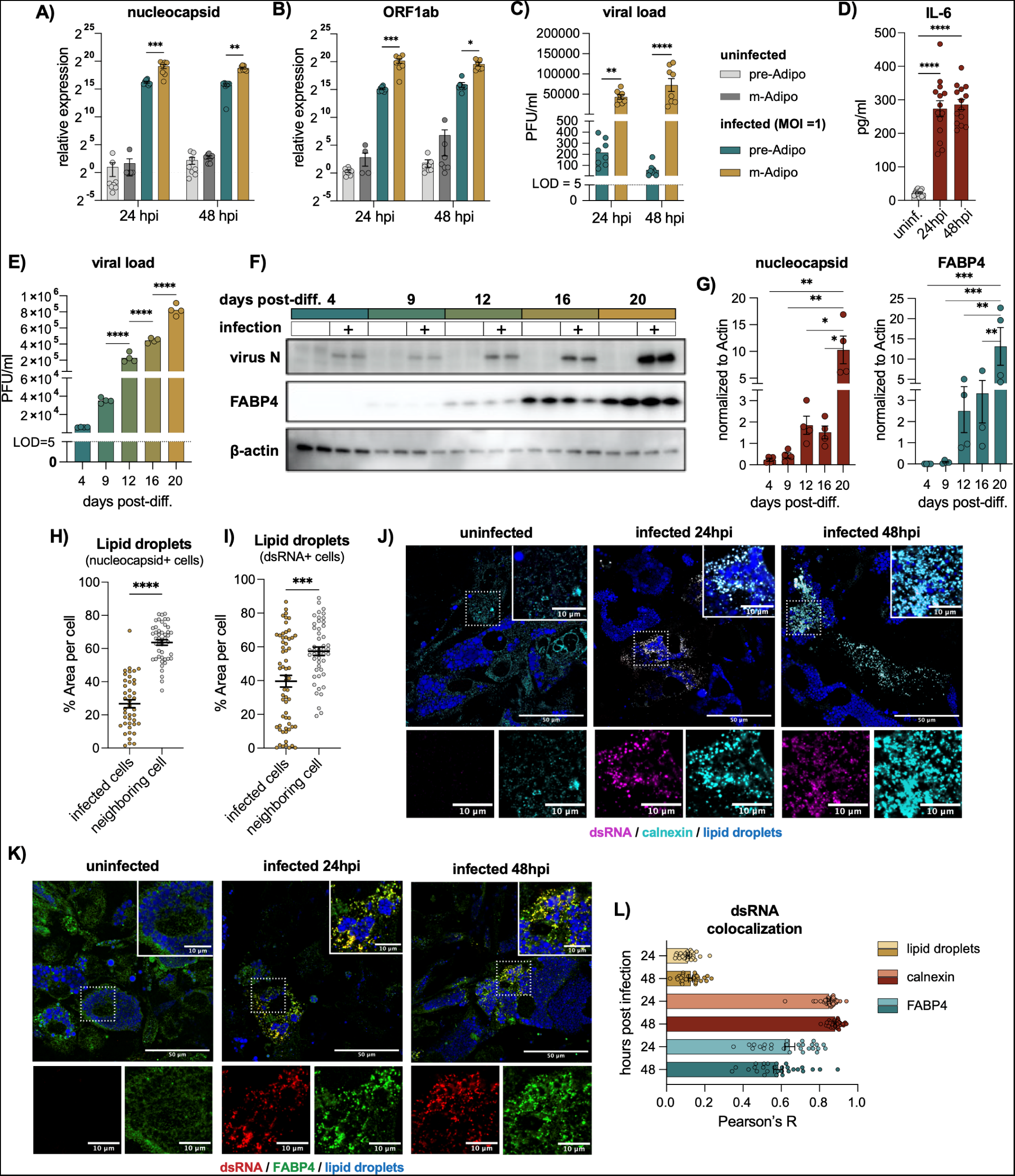
FABP4 colocalizes with SARS-CoV2 replication organelles in human adipocyte cell lines. A to D) hTERT pre-adipocytes and mature adipocytes infected with SARS-CoV2 (WA1/2020, MOI 1). A and B) relative expression of the virus A) genomic (nucleocapsid) and B) sub-genomic RNA (ORF1ab) normalized to β-actin. C) Viral loads measured from supernatant using plaque assay. Data is pooled from two independent experiments, (n=8). Statistical analysis was performed using two-way ANOVA. D) ELISA measuring IL-6 secretion from the supernatant of mature adipocytes in the presence or absence of virus infection (MOI 1). Data are pooled from three independent experiments (n=14). E to G) Adipocytes were infected at 4, 9, 12, 16, and 20 days, post-differentiation (MOI 1). Subsequent measurements were then performed 48 hours post infection. E) Viral loads measured by plaque assay. Data are representative of two independent experiments (n=4). D and E) Statistical analysis was performed using one-way ANOVA. F) Western blot of virus nucleocapsid, FABP4 and β-Actin from adipocyte lysates following infection. G) Quantification of the nucleocapsid and FABP4 band intensities normalized to β-actin. Data are representative of two independent experiments (n=4). Statistical analysis was done using two-way ANOVA. H and I) Percent area of lipid droplets of infected cells and neighboring cells, quantified based on fluorescence neutral lipid staining (Bodipy). Infected cells were identified by the H) nucleocapsid-positive staining, and I) dsRNA-positive staining. Data are pooled from two independent experiments (n=5) biological replicates, and statistical analysis were performed using standard t-test. J and K). Infected hTERT cells (MOI 1), (n=3) fixed 24 and 48 hours, post-infection. J) Cells stained for dsRNA (violet), calnexin (cyan) and lipid droplets (BODIPY, blue). Scale bar (50μm). K) dsRNA (red), FABP4 (green) and lipid droplets (blue) staining. Scale bar (50μm). J and K) Inlays represent magnified merged or single channel images of the areas indicated in J and K with a scale bar (10μm). L) Signal colocalization measure by Pearson’s R between dsRNA with calnexin, FABP4 and lipid droplets. Data are shown as the mean ± s.e.m.

To further examine the effect of infection on adipocytes’ lipid stores, we examined the size of lipid droplets (LDs) in infected cells, identified using either the nucleocapsid protein or the double stranded RNA (dsRNA), a product of viral replication detected at early stages of infection^1^ . We found that overall, the LDs of infected cells were significantly smaller compared to neighboring cells (Figures 2H and 2I). Notably, the size of the LDs in dsRNA-positive cells had a much wider variation compared to those in nucleocapsid-positive cells and showed a strong negative correlation with the dsRNA signal (Figures S2E and S2F). This correlation was clearly observable in our confocal images, where cells with fewer dsRNA puncti had larger LDs and vice versa (Figure S2G). These dsRNA puncti, were also positive for the ER membrane protein calnexin (Figure 2J and 2L), which condensed in infected cells in a pattern indicative of the virus DMVs ^32,33^. Overall, these results confirm that hTERT cells provide a good model to study SARS-CoV2 infection in human adipocytes and suggest that SARS-CoV2 replication is accelerated in cells of a higher lipid content, and that infection results in a gradual depletion of the cell’s lipid stores.

To contextualize the role of intracellular FABP4 during infection, we examined its expression, protein abundance, and secretion (Figures S2H–S2J). In this setting, FABP4 levels were similar across samples collected at 24 and 48 hours after infection. Fluorescence staining revealed a striking change in the spatial distribution of FABP4 within infected cells (Figures 2K–2L, and S2L). In dsRNA positive cells, FABP4 condensed into puncti that co-localized with the dsRNA signal (Figures 2K and 2L). Whereas in nucleocapsid positive cells, the FABP4 signal was distributed across the cytoplasm, showing a more heterogenous pattern of distribution (Figure S2L). Notably, both patterns were observed in neighboring cells, suggesting a dynamic change in FABP4 distribution at different stages of infection. These data show that FABP4 is recruited to the ROs and may play a role in virus replication.

### FABP4 deficiency impairs coronavirus replication

To understand the functional relevance of FABP4 in SARS-CoV2 infection, we utilized small molecule inhibitors of FABP4. For these experiments, we used two structurally different inhibitors targeting FABP4: BMS309403 ^34,35^ and CRE-14 ^36^, which represents a newly developed class of FABP inhibitors. We validated and confirmed the activity of these molecules in competing with FABP4 ligands using the Terbium-based time-resolved fluorescence energy transfer assay (TR-FRET), which confirmed that both BMS309403 and CRE-14 were able to block FABP4’s ability to bind its lipid ligand (BODIPY FL C12) (Figure S3C). BMS309403 has been reported to bind to FABP4 with a K_D_ of 552 nM ^35^. Using a micro-scale thermophoresis (MST) assay, we determined that CRE-14 binds FABP4 with a K_D_ = 954nM (Figures S3A and S3B).

Mature human adipocytes were treated with FABP4 inhibitors following SARS-CoV2 infection (MOI: 0.1 or 1). This inhibition significantly reduced virus nucleocapsid RNA and protein levels and resulted in a remarkable reduction in the virus titers (Figures 3A–3D, S3E and S3F). These results were further confirmed using confocal microscopy (Figures 3F and 3G). Moreover, FABP4 inhibition resulted in a significant reduction in IL-6 secretion 48 hours post-infection (Figure 3E), indicating an ameliorated inflammatory state. We then infected mature adipocytes with a higher viral dose (MOI: 3) to increase the incidence of dsRNA-positive cells, and thereby enable a better quantitative characterization of the dsRNA puncti. Interestingly, the dsRNA puncti in the FABP4-inhibitor treated samples appeared scattered across the cytoplasm failing to form coherent clusters (Figure 3H), indicating a disruption of DMV formation in which they are packed. A quantitative evaluation of the dsRNA positive signal in the confocal images confirmed this observation, revealing a significant reduction in the percentage of dsRNA-positive area and mean fluorescence intensity in infected cells following inhibitor treatment (Figure 3I).

**Figure 3:**
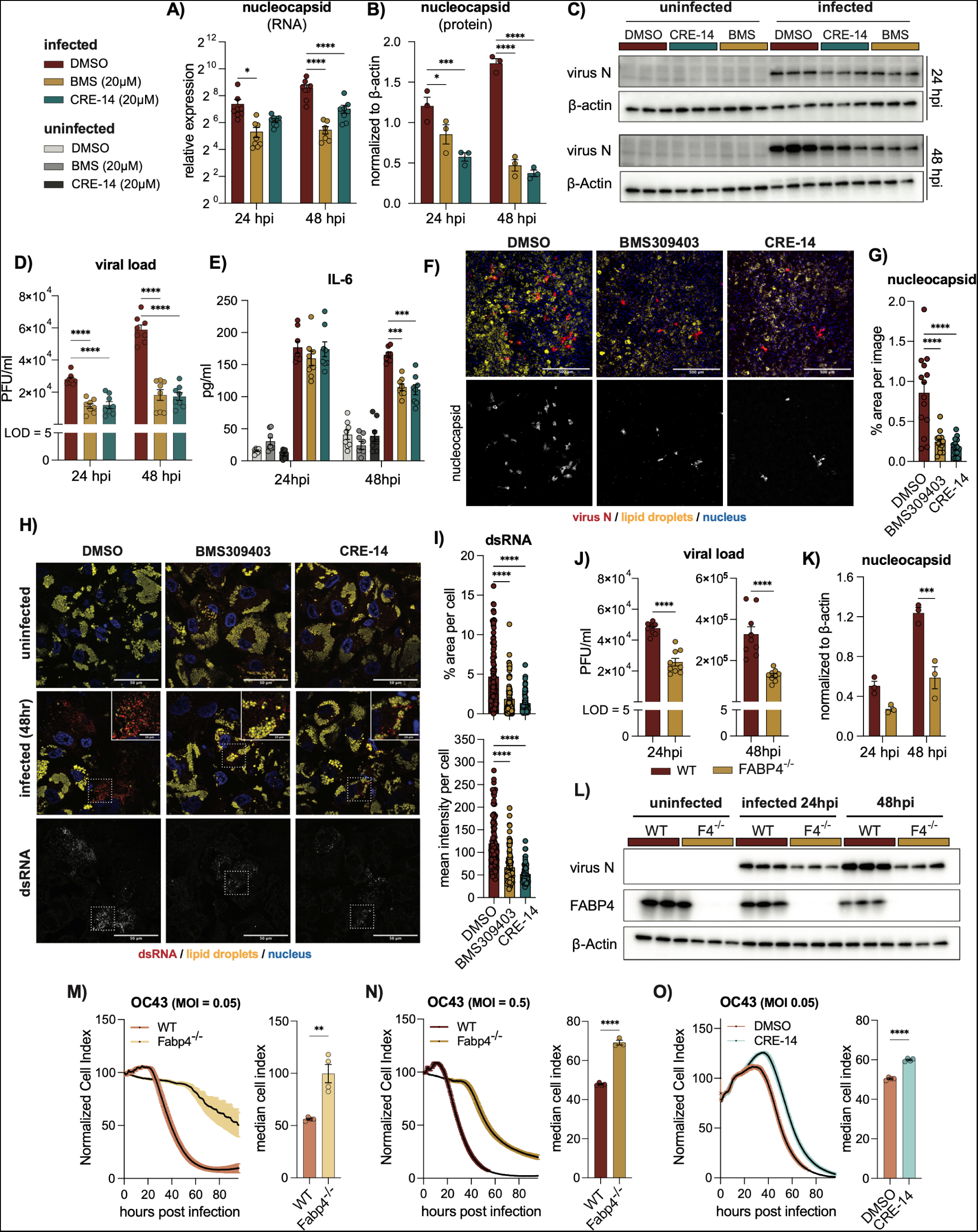
FABP4 deficiency reduces virus titers and disrupts replication organelles formation in adipocytes. A – G) SARS-CoV2-infected mature adipocytes (MOI:1), treated with either DMSO or FABP4 inhibitors, BMS309403 (20μM) or CRE-14 (20μM). A) Relative RNA expression of SARS-CoV2 nucleocapsid, normalized to β-actin. Data pooled from two independent experiments, (n=8). B) Western blot quantification evaluating nucleocapsid band intensity normalized to β-actin. Data are representative of three independent experiments (n=3). C) Western blot of nucleocapsid protein measured from cell lysates. D) Viral load measured by plaque assay. Pooled from two independent experiments, (n=8), representative of four independent experiments. E) ELISA measurement showing IL-6 secretion from infected adipocytes treated with FABP4 inhibitors. Data are pooled from two independent experiments, (n=8). A, B, D and C) Statistical analysis were performed using two-way ANOVA. F) Confocal images of adipocytes infected with SARS-CoV2 (MOI 1) then fixed 48 hours post-infection and stained for virus nucleocapsid (red), lipid droplets (BODIPY, yellow), and nuclei (DAPI, blue). (n=3) biological replicates. Scale bar (500μm). G) Percentage nucleocapsid positive area per image, with an average of 4-5 images per sample. H) Adipocytes infected with (MOI 3) and fixed 48 hours post-infection, then stained for dsRNA (red), lipid droplets (BODIPY, yellow) and nuclei (DAPI, blue). (n=3) Scale bar (50μm). Inlays represent magnified images of the indicated area with a scale bar of (10 μm). I) Percentages of dsRNA positive area and mean fluorescence intensity per cell. (n=3) biological replicates. Statistical analysis was performed using one-way ANOVA. J to L) Infected wild type (WT) and FABP4- deficient human adipocytes (MOI 1). J) Viral load measured by plaque assay from supernatants collected 24 and 48 hours post infected. Data are pooled from three independent experiments, (n=9). Statistical analysis was done using standard t-test. K) Quantification of nucleocapsid western blot band intensity normalized to β-actin. Data are representative of two independent experiments, (n=3). L) Western blot of nucleocapsid, FABP4 and β-actin proteins measured from cell lysates. M to O) Real time electric impedance traces and their corresponding median cell index (hours) of M and N) wild type and FABP4 knockout mouse pre-adipocytes infected with coronavirus OC43 at indicated MOIs. O) MRC5 cells infected with OC43 and treated with either DMSO or CRE-14. Statistical analyses were done using standard t-test (n=3). Data are shown as the mean ± s.e.m.

To further confirm the relevance of FABP4 in SARS-CoV2 replication in a genetic model, we generated FABP4-deficient human adipocyte cell lines through CRISPR-mediated deletion. We validated that this methodology was successful in eliminating FABP4 in these adipocytes, as shown by western blot analysis (Figure 3L). In these cells, viral titers and nucleocapsid protein abundance, measured at two different time points, were significantly reduced in the absence of FABP4 (Figures 3J – 3K). We also generated a second human adipocyte cell line expressing FABP4-targeting shRNA to achieve long-term suppression of FABP4 and observed a similar reduction in viral infection (Figure S3G).

These observations in multiple chemical and genetic models demonstrated the critical importance of FABP4 for SARS-CoV2 replication, likely through its engagement with the DMVs in the virus ROs. We next asked whether the effect of FABP4 on viral infection also applies to other common cold coronavirus infections. We examined common cold coronavirus (HCoV-OC43) infection using a Real-Time Cell Electronic Sensing assay (RTCES) to measure cell viability and cytopathic effects (CPE) in wild type and FABP4-deficient mouse pre-adipocytes (Figures 3M and 3N). We observed a significant reduction and delay in virus-induced CPE in the absence of FABP4. Similar results were also obtained in human lung fibroblasts (MRC5) following chemical inhibition of FABP4 (Figure 3O). Taken together, these results indicate that the effect of FABP4 may be broader and apply to multiple coronavirus infections.

### FABP4 facilitates virus replication in bronchial epithelial cells

Having established FABP4’s significance in SARS-CoV2 adipocyte infection, we investigated its role in human bronchial epithelial (HBE) cells to understand whether it is engaged early on during respiratory tract infection ^37^. HBE135-E6E7 cells were infected with high titers (MOI: 5) of various isolates of SARS-CoV2 representing the alpha, delta, omicron, and Eris variants (Ank1, Ank-Dlt1, and Ank-Omicron GKS, Ank-Eris respectively). Following infection, cells were treated with two doses of both FABP4 inhibitors, and their viral titers were monitored over four days (Figures 4A–4D). In all tested conditions, FABP4 inhibition resulted in a marked reduction of virus titers examined over the time course of 96 hours. Next, we used transmission electron microscopy (TEM) to examine the consequence of FABP4 inhibition on ROs morphology. Infected HBE cells (MOI:1) were treated with the FABP4 inhibitor (CRE-14) then fixed 48 hours post-infection. TEM imaging revealed a marked reduction in the size of DMVs following FABP4 inhibitor treatment (Figures 4SA and 4SC).

**Figure 4:**
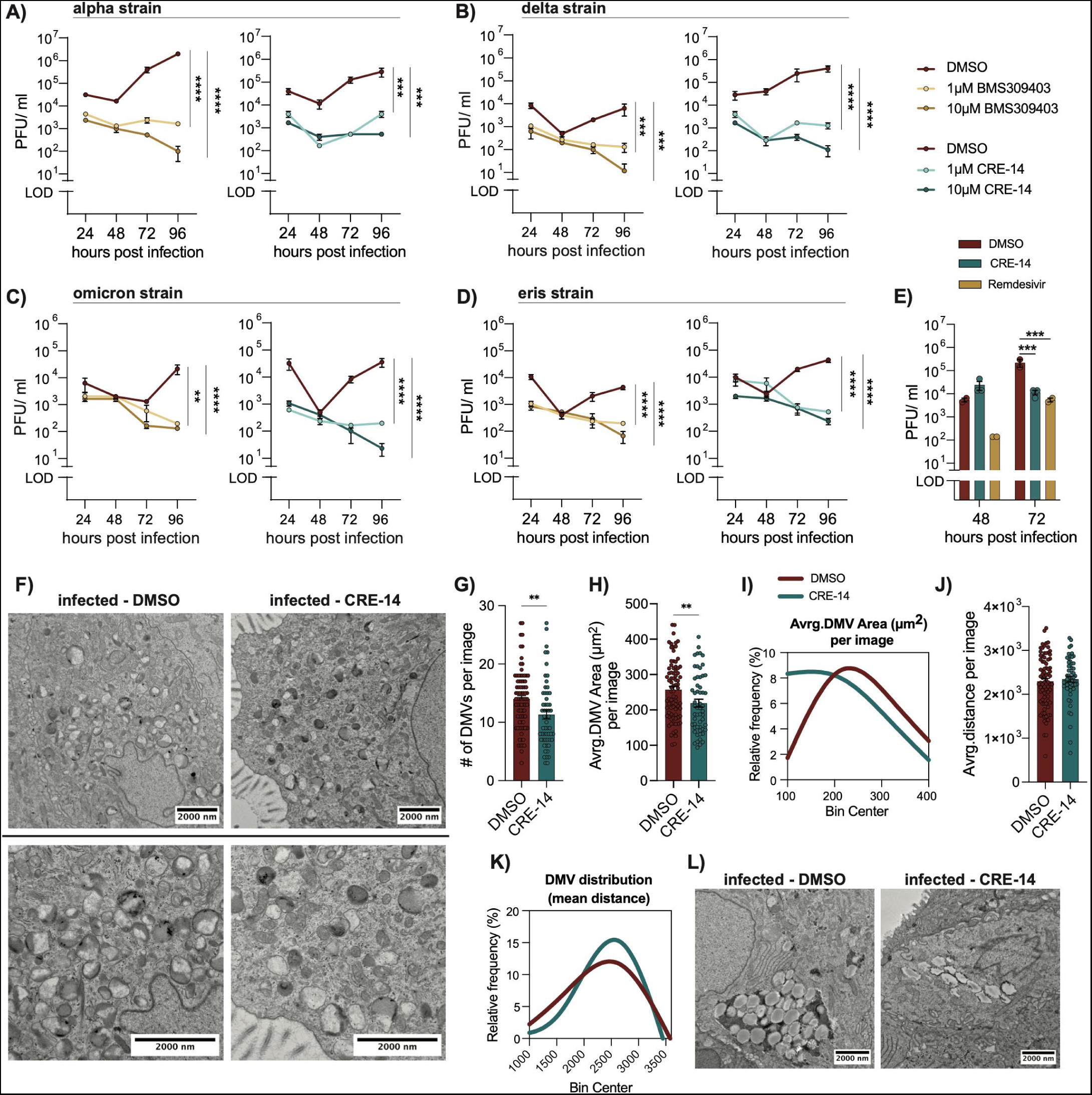
Inhibition of FABP4 in bronchial epithelial cells disrupts virus replication organelles. A to D) Viral load measured from the supernatant of human bronchoepithelial (HBE135-E6E7) cells infected with SARS-CoV2 (MOI: 5) A) alpha strain (Ank1), and B) delta strain (Ank-Dlt1) C) omicron strain (Ank-OmicronGKS) D) eris strain (Ank-Eris) and treated with BMS309403 or CRE-14 at the indicated doses. Data are representative of two independent experiments, (n=3). Statistical analysis was performed using two-way ANOVA. E) Virus load measured from the apical wash of infected 3D airway epithelium cultures (IDF0571/2020, MOI 0.1), treated with either CRE-14 (10μM) or Remdesivir (5μM). Statistical analysis was performed using two-way ANOVA. (n=3). F to L) 3D reconstructed airway epithelium cultures were infected apically with 10^5^ PFU of SARS-CoV2 (strain WA1/2020) then treated with either DMSO or CRE-14 through the basal layer. 24 hours after infection, the cells were fixed for EM and 3 sections per sample (n=2) were analyzed. F and L) Representative TEM images showing F) virus double-membrane vesicles (DMVs) and L) lipid droplets in control (DMSO) and inhibitor treated samples. G) Number of double membrane vehicles and H) their average area per image. (n=2) biological replicates with 18-45 images per sample taken at an 8000X magnification. I) Frequency distribution of DMV area shown as a Fit Spline. J) The mean distance between DMVs within each image, calculated from the X, Y coordinates of each DMV and K) the percent frequency distribution shown as a fit spline. Statistical analyses were done using standard t-test. Data are shown as the mean ± s.e.m.

We also utilized reconstructed airway epithelium organoids that were apically infected with SARS-CoV2 and treated with 10μM CRE-14 or 5μM of the antiviral Remdesivir through the basal layer. Viral titers measured from the apical wash showed a reduction following inhibitor treatment 72 hours post-infection that, at the higher dose, matched the titers of the Remdesivir treated group (Figure 4E). We then examined the morphology and distribution of DMVs in the vehicle or CRE-14 treated organoids and observed that in the inhibitor-treated group DMVs were reduced in both size and numbers (Figures 4F–4I) and appeared more dispersed, failing to form the tight clusters observed in untreated controls (Figure. 4J and 4K). Clusters of LDs were observed in infected samples and were nearly absent in the inhibitor-treated group (Figure 4L). Importantly, we confirmed that exposure to FABP4 inhibitor in uninfected samples did not alter cellular or organelle morphology (Figure. S4B). We performed a similar analysis in samples treated with the BMS309403 compound, and though we observed similar results, the phenotype was less pronounced, and the compound treatment showed signs of toxicity in these organoids (Figures. S4D–S4F). Overall, these data confirm the importance of FABP4 for SARS-CoV2 replication through its influence on the ROs formation in multiple cellular targets. Its engagement with virus infection in airway epithelial cells and adipocytes, also increases its potential as a therapeutic target.

### FABP4 inhibition reduces viral titer and lung damage in infected hamsters

To examine the therapeutic potential of FABP4 inhibitor treatment in a preclinical model *in vivo*, we infected 12–14-week-old lean Syrian hamsters with SARS-CoV2 (Ank1, 100 TCID_50_) ^38^, and treated them for 6-days with a subcutaneous injection of 15 mg/kg FABP4 inhibitor BMS309403, CRE-14 or vehicle (Figures 5 and S5). The hamsters were monitored daily, and the experiments were terminated for virus titer measurements (right lung) and histological analysis (left lung). Inhibitor treatment significantly ameliorated the infection-associated weight loss (Figure 5A), and the viral titers were consistently decreased upon treatment with FABP4 inhibitor (Figure 5B).

**Figure 5:**
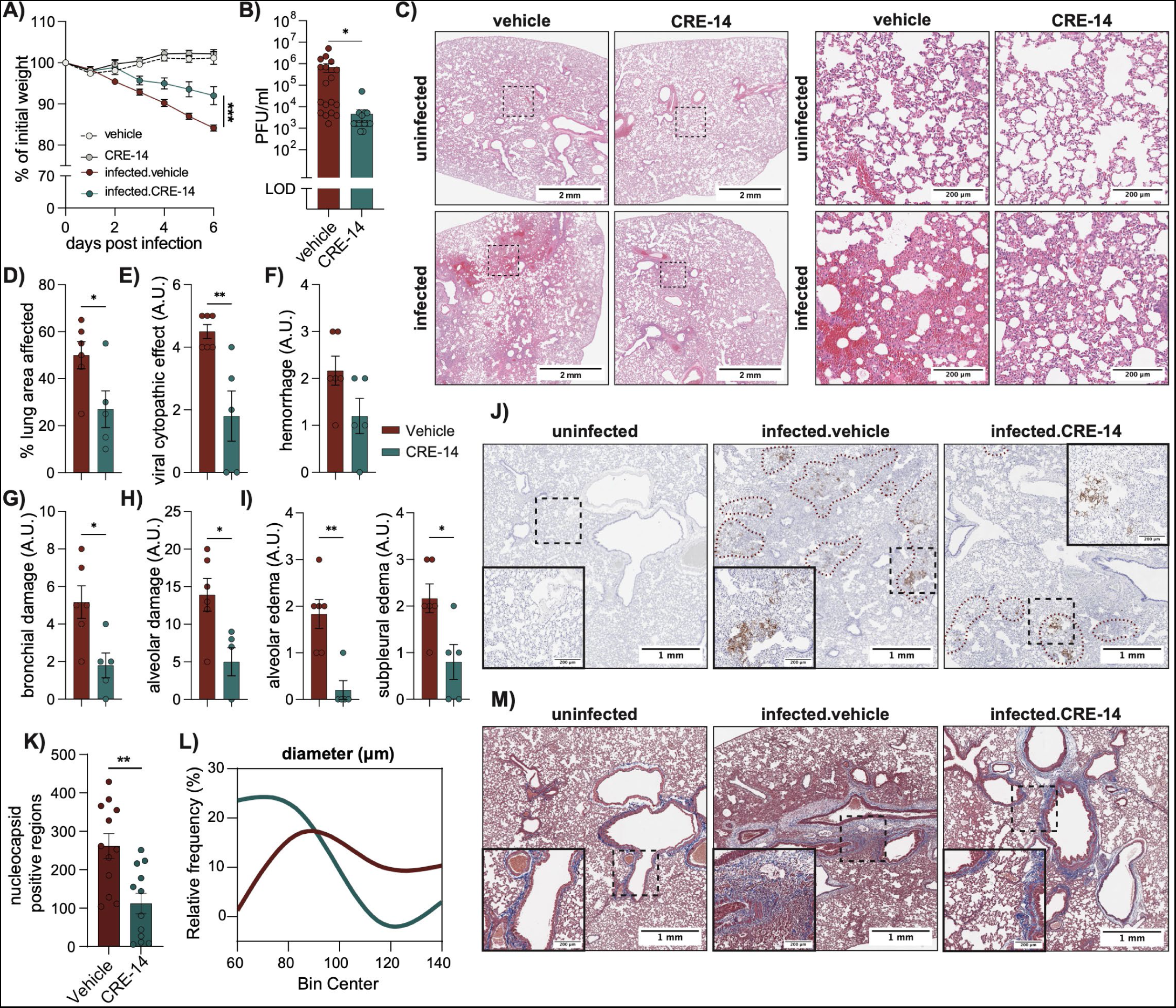
FABP4 blockade limits virus replication and ameliorates pathology in infected hamsters. Syrian hamsters were infected intranasally with SARS-CoV2 (Ank1, 100 TICD_50_) and treated daily with FABP4 inhibitor (15mg/kg, CRE-14) or vehicle. A) Percent of initial body weight. B) Lung viral titers pooled from three independent experiments. (n=18 and 17 for infected vehicle and CRE-14 respectively and n=6 for each of the uninfected groups). Statistical analysis was performed using standard t-test. C) Representative H&E staining of hamster lungs with or without inhibitor treatment. Scale bars (left: 2mm, right 200μm). D to I) Pathology evaluation of lung histology. Arbitrary units (A.U.) is derived from the pathology score. Further details are found in the extended data. Statistical analysis was performed using standard t-test. G) Bronchial damage data shows the combined pathology scores of the bronchial epithelial cell necrosis and presence of cellular debris in bronchi. H) Alveolar damage shows the combined pathology scores of alveolar epithelial cell necrosis, cellular debris in alveoli, hyaline membranes, fibrin deposition and alveolar emphysema. J) Representative IHC of SARS-CoV2 nucleocapsid staining of hamster lungs. K) Number of nucleocapsid positive regions and L) the frequency distribution of their diameter, as quantified from the lung IHC represented in J. M) Representative images showing Masson’s trichrome staining of hamster lungs. Data shows the mean ± s.e.m.

We further examined the effect of FABP4 inhibition on lung pathology and observed a notable reduction in lung damage (Figures 5C and S5D). A careful evaluation of these samples performed by an independent pathologist showed a reduction in various clinical aspects associated with viral infections, including lung damage within bronchial and alveolar areas, as well as alveolar and subpleural edema (Figures 5D–5I). Moreover, using the same sample set, we performed immunohistochemical staining for the SARS-CoV2 nucleocapsid protein, which revealed that nucleocapsid positive areas were reduced both in number and in size after FABP4- inhibitor treatment (Figures 5J–5L). Trichrome staining showed increased collagen deposition in the lungs following infection which was reduced with inhibitor treatment (Figure 5M and S5E). Hamster treatment with BMS309403 resulted in similar but less pronounced results (Figures S5F and S5G). This might be explained in part by the higher plasma exposure and better pharmacokinetics of CRE-14 as assessed in mice (Figures. S5A–S5C).

## Discussion

In this study, we identify FABP4 as a critical host factor for SARS-CoV-2 infection and pathogenesis. We establish FABP4’s importance in two separate human COVID-19 cohorts and demonstrate, through *in vitro* studies, that targeting FABP4 effectively reduces viral titers in multiple cell types and viral strains. Additionally, the reduced cytopathic effects we observed in the absence of FABP4 in cells infected with the common cold coronavirus OC43, indicates that targeting FABP4 may be effective across various coronavirus infections. We also demonstrate that FABP4 inhibitor treatment of infected Syrian hamsters decreased viral loads and alleviated the infection-associated immunopathology.

Remodeling of the host intracellular membranes into virus replication organelles (ROs) is a strategy used by coronaviruses and other positive stranded RNA (+RNA) viruses, to support their propagation and secretion and provide an environment shielded from immune recognition ^2^. In the case of coronaviruses, these compartments consist of DMVs, spherules, and zippering ER membranes. In this study, we observed a striking recruitment of FABP4 to ROs in infected cells. The FABP4 signal, which is usually diffused throughout the cytoplasm, condensed into puncti colocalized with dsRNA signals, pointing to their recruitment to DMVs. Examination of infected adipocytes and bronchial epithelial cells with confocal microscopy and TEM revealed that the DMVs in cells where FABP4 was inhibited were smaller and more dispersed. We suggest that this loss of a spatial organization of the DMVs is what resulted in the reduced viral replication capacity, evident by the lower virus nucleocapsid RNA and protein, and overall viral titers. Investigating the exact role that FABP4 serves within the ROs is an exciting avenue to explore in future research, with potential implications for novel viral intervention strategies. Previous studies have shown that inhibiting *de novo* lipogenesis (DNL) or disrupting the contact sites between LDs and the RO zippering ERs diminishes virus titers^39,40^. We anticipate that FABP4 might be an important upstream regulator of DNL ^18,41^ in the early phase of infection, that promotes the accumulation of LDs and their later utilization by facilitating lipid trafficking to the membranes of viral ROs ^17^. Interestingly, our examination of SARS-CoV-2 infection dynamics in human adipocytes revealed that adipocyte maturation significantly enhances viral replication capacity. Though this could be attributed to increased lipid accumulation, the maturation process is also coupled to a substantial increase in FABP4 abundance. We observed a rapid depletion of lipid droplets in infected adipocytes that is proportional to the density of their dsRNA puncta indicating that LDs are utilized during RO biogenesis and may be essential in facilitating it. This observation aligns with the work published by Ricciardi *et al*.^40^ showing that cells transfected with virus non-structural proteins (nsp 3,4 and 6) to form RO-like structures (ROLS) also accumulate, then consume LDs during the process of ROLS formation. A recent study reported an increase in the LD area of infected Simpson-Golabi-Behmel syndrome (SGB)-derived adipocytes^42^. This might be a result of the timepoint in which this analysis was made in addition to variations in dynamics of cellular responses to infection across the hTERT and SGB-derived adipocytes. These observations might provide one possible explanation of how obesity can exacerbate disease severity in COVID-19, both due to the higher lipid content in adipocytes and the ectopic lipid deposition at other organs associated with obesity ^41^.

FABP4 has a marked impact on the metabolic dysregulation associated with obesity and aging ^43^. Its increase in obese mice and humans correlates positively with cardiometabolic pathologies and morbidity ^14^. These correlations were clearly reflected in our patient cohorts and were associated with more severe COVID-19 disease. Furthermore, prior GWAS studies reported that humans carrying a low-expression variant of FABP4 exhibit significant protection against the development of type 2 diabetes and cardiometabolic disease ^44–46^. Consequently, the impact of targeting FABP4 on the overall pathogenesis of COVID-19, and the protection that such interventions may offer to individuals with underlying metabolic conditions, may not be limited to the reduction in viral titers. FABP4 can also influence the inflammatory and metabolic milieu both within the local environment of the lungs, and systemically through its action as a secreted hormone. In support of this possibility, we have observed a marked induction of IL-6 production in SARS-CoV2-infected adipocytes and significant loss of this response in the absence of FABP4. IL-6 is the most commonly used clinical risk biomarker and its potential emergence from adipose tissue is an intriguing prospect of how adipocyte infection may be a critical determinant of disease course.

In animal models of allergic asthma, FABP4 expression is increased in lung epithelial and endothelial cells, and its genetic deletion improved airway function and attenuates inflammation and immune cell recruitment ^16,47^. Similar mechanisms could explain the reduced lung damage and collagen deposition that we observed in our inhibitor-treated, SARS-CoV2-infected hamsters. Moreover, we’ve shown in previous studies that antibody-mediated neutralization of FABP4 improves metabolic health in obese mice by lowering hepatic glucose production, improving insulin sensitivity, and reducing hepatic steatosis^13,48^. These effects may also counteract the COVID-19-associated hyperglycemia which is also attributed to increased hepatic glucose production and insulin resistance ^26,49^. While this requires further investigation, a thorough examination of FABP4’s engagement in these pathways could have important implications for mitigating the risk of developing diabetes or sustaining long-term respiratory symptoms, both documented amongst the many symptoms of long-COVID ^50,51^. Such investigations would benefit from the use of mouse models, for which we have a much more expansive toolkit of reagents, genetic, and pharmacological means of studying FABP4 biology, compared to hamster models. Thus, in future research, we hope to expand on of the findings of this study using mouse-adapted coronavirus strains^52,53^.

FABP4 protein levels in the circulation are markedly elevated in humans with severe to critical clinical manifestations following SARS-CoV2 infection, which points to the possibility of infected adipocytes producing higher levels of the protein. However, in the isolated cultured adipocytes, we did not detect an increase in FABP4 secretion upon infection. It is possible that adipocytes in their native tissue environment respond differently to infection, or that FABP4 secretion *in vivo* is elicited as a response to more complex systemic interactions, particularly those associated with the pathophysiology of severe COVID-19. This would explain why individuals with mild to moderate disease have significantly lower concentrations of FABP4. It is also possible that the source of FABP4 in circulation may be endothelial cells as they do provide a substantial proportion of circulating FABP4, especially at the baseline state ^54^. This would be quite interesting to explore in the future in the cell type-specific FABP4-deletion mouse models.

In summary, our works demonstrates a critical contribution of FABP4 to SARS-CoV2 infection as a host factor, and highlights FABP4 as a therapeutic target acting both as an antiviral and a modulator of cardio-pulmonary and metabolic fitness.

## Acknowledgments

We thank all members of the Hotamisligil Laboratory, past and present, for their scientific support and input during the development of this project. We also thank Drs. Phyllis Kanki, and Don Hamel for their support in facilitating our work in the BSL-3 facility at Harvard Chan School, Dr. Lynn Enquist of Princeton University for his valuable feedback, Dr. Pinar Firat from Koç University and Dr. Nur Urer from the Ministry of Health Yedikule State Hospital for help with human histological samples. Electron microscopy image analysis was conducted with assistance from the Image Analysis Collaboratory at Harvard Medical School (https://iac.hms.harvard.edu/).

## Author contributions

HB designed and performed the *in vitro* and cell-based experiments and analyzed the data, performed the histology, fluorescence, and electron microscopy image analysis, prepared the figures, and wrote and revised the manuscript. EK designed, supervised, and performed experiments relating to the CRE-14 compound and OC43 virus infection, interpreted results, revised the manuscript, and contributed to the conception of the project. GT and FB provided intellectual contributions and coordinated collaborations. HK designed and synthesized the CRE-14 compound and provided intellectual contributions and contributed to the conception of the project. LM generated the human FABP4 knockout cell lines and together with NB and ESK performed sample analysis. GYL provided intellectual insight and assistance in generating the human FABP4 knockout cell lines. AH and ZFK performed *in vivo* hamster experiments. NF, and BG performed *in vitro* assays and analyzed data. SO, AC, OOC, and FS performed histological sample preparations and analyzed the data. AA Established the RTCES assay protocols and analyzed data. EY, IE and AGE maintained the animal colonies and designed and prepared animal cohorts and supervised animal experiments. FC and ÖE conducted all human studies and performed the FABP4 measurements in human samples. HL and XL performed the statistical analysis for the human data. AÖ planned, supervised, and conducted all hamster infection experiments and analysis of the results and revised the manuscript. GSH conceived, supervised and supported the project, designed experiments, interpreted results and planned and revised the manuscript.

## Supplementary Materials

Materials and Methods:

Figs. S1 to S5

