## Supplementary Information for "FABP4 as a Therapeutic Host Target Controlling SARS-CoV2 Infection"

**The PDF file includes:**

- Materials and Methods
- Figs. S1 to S5

### Materials and Methods

Cell lines:

human Telomerase Reverse Transcriptase (hTERT) pre-adipocytes [1], were maintained in DMEM/high glucose media (ThermoFisher Scientific, cat.no. 11965118) with 10% FBS (Atlanta Biologicals, cat.no. S11550), 1% penicillin streptomycin (Lonza, cat.no. 17-603E). Cells were differentiated two days after reaching confluence using DMEM/high glucose with 2% FBS, 1% penicillin streptomycin, human insulin (0.5µM, Sigma Aldrich, cat.no. I9278), biotin (33µM, Sigma Aldrich, cat.no. B4639), panthothenate (17µM, Sigma Aldrich, cat.no. P-5155), dexamethasone (0.1µM, Sigma Aldrich, cat.no. D-1756), Triiodothyronin (2nM, Sigma Aldrich, cat.no. T-6397), IBMX (500µM, Sigma Aldrich, cat.no. I-5879), Indomethacin (30µM, Sigma Aldrich, cat.no. I8280). Cells were maintained in differentiation media for 18-20 days prior to the start of infection.

Wildtype and *Fabp4* knockout 3T3-L1 mouse adipocyte cell lines [2] were maintained in DMEM/high glucose 10% BCS (Hycline, cat.no. 16777-206), and 1% penicillin streptomycin, then differentiated two days after reaching confluence with DMEM/high glucose, 10% FBS, 1% penicillin streptomycin, 5mg/ml insulin, 500mM IBMX, and 2mM rosiglitazone (Cyman, cat.no. 71740). Cells were kept in differentiation media for two days then switched to growth media with 1mg/ml insulin.

Vero-E6 cells (ATCC, CRL-1586) were cultured in DMEM/high glucose with 10%FBS, 1% penicillin streptomycin, 1% HEPES (ThermoFisher Scientific, cat.no. 15630080). HBE135-E6E7 human bronchoepithelial cells (ATCC, CRL-2741), were cultured in DMEM/high glucose with 10%FBS, 1% penicillin streptomycin. Human MRC5 lung fibroblasts (ATCC, CCL-171) were cultured in DMEM/high glucose with 10% FBS, 4mM L-glutamine, and 1% penicillin streptomycin. Airway epithelium organoids (MATTEK, cat. no. AIR-100, AIR-112) were cultured following manufacturer's instructions. Viral titers measurements in airway epithelium organoids were performed by Epithelix Sarl, France (www.epithelix.com), using MucilAir™ human 3D tissue from airways and lung surgical pieces, following manufacturer's instruction. FBS used in all cellular experiments was heat inactivated.

Viruses:

SARS-CoV2 (USA-WA1/2020) isolate seed stock was acquired from BEI resources (Cat.no. NR-52281) and propagated in-house in Vero-E6 cells. SARS-CoV2 (Ank1) was isolated from nasopharyngeal swabs of patients with COVID-19-related respiratory illness in January 2020, Ankara, Turkey, and shown to be an alpha variant of SARS-CoV2 (B.1.1.7) [3], (GenBank Acc. No: MT478019). The SARS-CoV2 Delta, Omicron and Eris variants were isolated from patient nasopharyngeal swabs in September 2021, April 2022, and May 2023 respectively. [Ank-Dlt1 (B.1.617.2) (GenBank Acc. No: OM295705), and Ank-Omicron GKS (BA.1.17) (GenBank Acc. No: OR529199)]. The variant identification was done after genome wide sequence analysis following two-step plaque purification for the Delta and Omicron variants. The Eris variant was identified through partial sequencing of the spike coding region. SARS-CoV-2 BetaCoV/France/IDF0571/2020 strain was used in the MucilAir™ 3D human airway epithelial cell infections.

Human β-coronavirus OC43 (ATCC, VR-1558), was propagated in MRC5 cells. All infections were performed using growth media (corresponding to the infected cell) with 2% heat inactivated FBS. Target cells were incubated with a low volume of virus-containing media for 1 hour, with gentle shaking every 15 minutes. After infection, cells were washed with PBS before

adding fresh growth media. 3D airway epithelium cultures were infected through the apical layer after it was washed with TEER buffer (MATTEK, TEER-BUFFER, PBS with $Mg^{2+}$ and $Ca^{2+}$). The apical layer was washed again following infection, and liquid was removed.
All *in vitro* experiments involving SARS-CoV2 were performed in Biosafety level 3 facilities at the Department of Immunology and Infectious Diseases at Harvard T.H Chan School of Public Health, VirNext (France) and Faculty of Veterinary Medicine, Ankara University, following the approved standard operating procedures.

Human data:
Ethics approval was obtained from Koç University Institutional Review Board (IRB) for Clinical Research (Approval number: 2020.136.IRB1.026). Lung biopsies were acquired between June-September of 2020 from a control subject and 3 COVID-19 patients: patient 1 (age 61) underwent a biopsy for squamous cell carcinoma, and patients 2 and 3 (ages 46 and 64) underwent biopsy for hemoptysis symptoms. All patients tested positive for COIVD-19 on the day of biopsy. The control lung biopsy was acquired from an 85-year-old male patient with squamous cell carcinoma, collect in 2021, with a negative COVID-19 test. Serum samples from COVID-19 patients and healthy controls were collected at the Koç University Medical School Hospital.

Hamster infection:
Specific Pathogen Free (SPF) Syrian Hamsters (Mesocricetus auratus, 12-14 weeks old, males and females of 120 g average body weight at time of infection) were purchased from Janvier laboratories, France (cat. name GOLDHAMSTER, strain RjHan:AURA), and then maintained in the SPF animal facility at the Biotechnology Institute, Ankara University, accredited by the Ministry of Agriculture and Forestry, Turkey. Hamsters were housed under a 12:12 light: dark cycle and fed ad libitum.
*In vivo* SARS-CoV2 studies were conducted in Animal Biosafety level 3 laboratories in Faculty of Veterinary Medicine, Ankara University. Humane endpoint scores were considered and multiple observations per day were conducted to confirm the animals' welfare. Hamsters were anesthetized using a combination of 100mg/kg Ketamine and 7mg/kg xylasine, then infected intranasally with SARS-CoV2 (Ank1) dissolved in serum-free DMEM. Inhibitor treatments were administered daily by subcutaneous injection, starting simultaneously with infection. At the experiment endpoint (on day 6 post-infection), hamsters were euthanized after anesthesia. All animal experiments were performed with the official permission of the Local Ethical Committee for Animal Experiments, Ankara University (06/May/2020, 2020-08-66 and/09/2021, 2021-16-150). All animal samplings were conducted according to the national regulations on the operation and procedure of animal experiment ethics committees (Regulation Nr: 26220, Date: 09.7.2006).

FABP4 inhibitors:
CRE-14 was manufactured and provided by Crescenta Biosciences Inc. as described in the US Patent 17/566692 published on July 6, 2023 [4]. The CRE-14 interaction with FABP4 was evaluated using a TR-FRET-based ligand displacement assays and micro-scale thermophoresis assay (MST). The ligand displacement assay was performed by Cepter Biopartners, New Jersey, (https://www.cepterbiopartners.com/), using a Terbium (Tb)-based time-resolved fluorescence energy transfer (TR-FRET) assay. Briefly, CRE-14 and BODIPY FL C12 were prepared at a

concentration of 1.085 mM and 4.2 µM, respectively, in DMSO. 1.2 µL of each compound or DMSO (vehicle control) and 1.2 µL of BODIPY FL C12 were added into the wells of a 384-well black polypropylene plate. His6-FABP4 and Tb anti-His6 antibody were prepared in the assay buffer (25 mM Tris/HCl, pH 7.4, 0.4 mg/ml ɣ-globulins, 0.010% NP-40, 1 mM DTT) at a concentration of 83 nM and 49.6 nM, respectively. The protein and antibody solutions were then mixed at a ratio of 34:7 (v/v) and incubated on ice for 30 minutes. The assay was initiated by adding 41 µL of the resulting protein/antibody solution into the wells containing the compounds and BODIPY FL C12. The plate was centrifuged and incubated at room temperature for 10 min. The TR-FRET signals were detected using an EnVision Multilabel plate reader (PerkinElmer; TB excitation 320 nm, BODIPY FL C12 emission 520 nm; TB emission 615 nm). Relative fluorescence ratio (520nmx10,000/615nm) was used to calculate the compound mediated inhibition of BODIPY C12 FL fatty acid binding to FABP4. The same procedure was performed with the BMS309403 compound. The MST assay was performed by 2bind GmbH, Germany (https://2bind.com/). Recombinant human FABP4 produced *E. coli* was labeled with a fluorescent dye NT-650-NHS 2nd generation fluorescent dye (Nanotemper) in the labeling buffer (1 x PBS pH 7.4, 1 mM TCEP, 2.5% DMSO) containing 10 µM of the recombinant FABP4 and 30 µM of the dye at 25°C. Following a 30-minute incubation the solution was passed through PD Minitrap™ G-25 columns to eliminate the excess dye. Serial dilutions of CRE-14 were prepared in the assay buffer and mixed with the labeled FABP4 in a final volume of 10 µl of the assay buffer (1 x PBS pH 7.4, 1 mM TCEP, 0.05% Pluronic F-127, 2% DMSO). The solution was filled into premium-coated glass capillaries (NantoTemper) and analyzed on a Monolith NT.115 Pico machine (red-pico; NanoTemper) at 25°C. The final concentration of the labeled FABP4 was 10nM and the CRE-14 concentration ranged between 20 µM to 610 pM (15X 2-fold dilutions of the highest concentration of the compound). The binding affinity constants ($K_D$) for FABP4 were estimated by fitting the dose-response curve using non-linear regression using GraphPad Prism software.

BMS309403 (MedKoo, cat.no. 524464) is a highly selective FABP4 inhibitor. Further information on this compound was previously reported [5]. For cellular experiments FABP4 inhibitors (BMS309403, CRE-14) were dissolved in sterile DMSO (Cell Signaling Technology, cat.no. 12611S), to create a stock solution of 10mM, which were aliquoted and stored at -20°C. Remdesivir (MedChemExpress cat. No. HY-104077) was diluted in DMSO and used at 5 µM. The impact of CRE-14 on cell viability was assessed by incubating MRC5 cells with increasing concentrations of the compound using media containing either 2% or 10% FBS. 72 hours after incubation, cell viability was measured using ONE-Glo™ Luciferase assay kit (Promega, cat.no. E6110). Prior to cell treatments, a solution of the desired concentration of the inhibitors were created by dissolving the inhibitors in the respective cell growth mediums, containing 2% heat inactivated FBS or 1mg/ml bovine serum albumin (Carl Roth, cat.no. 9048-46-8). Inhibitor treatments were initiated 1-hour after cell incubation with infection media.

For *in vivo* treatment, inhibitors were dissolved in 0.5% hydroxypropyl methylcellulose (HPMC) (Merck, cat.no.H7509) containing 1% Tween-80 (Sigma-Aldrich, cat.no. P1754) and the pH of the drug was adjusted to 8.0 before injection. A solution of (0.5% HPMC and 1%T-80, pH8.0) was used as vehicle. The Pharmacokinetics of BMS309403 and CRE-14 *in vivo* were assessed in C56BL/6J mice and Syrian hamsters following subcutaneous injection. Blood samples were collected at the indicated intervals and the compounds' concentration in the plasma were assessed by LC MS/MS.

Virus infectivity assays:
For plaque assay quantification of viral loads, ~$2x10^5$ cells/well of Vero-E6 cells were seeded in 12-well plates 24 hours prior to infection. 10-fold serial dilutions of the supernatant samples were created using DMEM/high glucose media with 2% heat inactivated FBS and 1%HEPES. After removing the Vero cell growth media, 200µl of samples were added and incubated for 1 hour with gentle shaking every 15 minutes. After incubation, an overlay of (1:1) growth media and 2% methylcellulose (Sigma Aldrich, cat.no. M0512) was added, and the cells were incubated at 37°C for approximately 72 hours. The overlay was then removed and 4% formaldehyde (Fisher Scientific, cat.no. BP531-500) in PBS was used to fix the cells for 20 minutes. Cells were then washed with PBS, and then stained using crystal violet solution to quantify plaques [2%(w/v) crystal violet (Sigma Aldrich, cat.no. C0775-25G), with 20% methanol in distilled water]. Viral titers measured from HBE cells, 3D reconstructed airway organoids and hamster lung viral titers were evaluated as TCID50 measurements (50% tissue culture infectious dose), then converted to plaque forming units (PFU= TCID50 x 0.7). Here cytopathic effects were continually observed under a light microscope for each time point, and the TCID50 was calculated using the Reed and Muench method. Lung viral loads in the hamster studies were measured from the inferior and post-caval lobes of the lungs after homogenization in DMEM.

CRISPR deletion and shRNA targeting FABP4:
Genetic deletion of *FABP4* in human adipocytes cells (hTERT) was performed using lentiviral CRISPR plasmids. The lentiviral CRISPR plasmids were generated by cloning single guide RNA targeting FABP4 or control sgRNA into lentiviral CRISPR backbone lentiCRISPRv2 (Addgene, cat.no. 52961). HEK293 cells (ATCC, cat.no. CRL-1573] were transfected with the lentiviral plasmids using Lipofectamine™ LTX Reagent with PLUS™ Reagent (ThermoFisher Scientific, cat.no. 15338100). Lentivirus particles were collected from the filtered supernatant and used to infect hTERT cells (at a 40-50% confluence) with the addition of 5µg/ml polybrene (EMD Millipore, cat.no. TR-1003-G). hTERT cells were re-plated from 6cm to 10cm dishes once they reached 80% confluence and were subjected to 1µg/ml puromycin (Sigma-Aldrich, cat.no. P4512) selection.
hTERT cell lines stably expressing shRNA targeting FABP4 or a scrambled control were generated using lentiviral shRNA plasmids (Origene cat.no. TL313105, and TR30021 respectively). HEK293 cells were transfected using Lipofectamine™ LTX Reagent with PLUS™ Reagent, and lentiviral particles were collected from filtered supernatant and used to infected hTERT cells with the addition of 8µg/ml Polybrene. After 24 hours, virus medium was removed, and a fresh growth media was added. After another 24 hours, hTERT cells were re-plated onto 10cm dishes and subjected to 3µg/ml puromycin selection. To validate the CRISPR knockout and shRNA knockdown of FABP4, cells were differentiated and FABP4 protein levels were examined at different stages of differentiation.
3T3-L1 mouse adipocytes lacking *Fapb4* were reported earlier by our laboratory [2], and were maintained and differentiated using the same protocol as the wildtype 3T3-L1 adipocytes.

RNA isolation and qPCR:
Qiazol reagent (Qiagen, cat.no. 79306) was used to inactivate SARS-CoV2 and collect RNA from cell lysates following the manufacturer's instructions. The RNA-containing aqueous phase was then added to an equal volume of 70% ethanol and RNA isolation was conducted using the NucleoSpin RNA isolation kit (Mecherey-Nagel, cat.no. 740955). cDNA synthesis was

conducted using the iScript cDNA synthesis kit (BioRab, cat.no. 1708897), and the qPCR was performed using Taqman reagents [ThermoFisher Scientific; SARS-CoV2 *nucleocapsid* (Vi07918637_s1), SARS-CoV2 *ORF1ab* (Vi07921935_s1), human *FABP4* (Hs01086177_m1), human *β-actin* (Hs01060665_g1)].
In hamster studies, virus RNA was extracted from lung homogenate of the superior right lobe in 1ml of Triazole Reagent (ThermoFisher Scientific, cat.no. 15596026). Virus genomic RNA for the spike encoding gene was measured using the previously described primer/probe set (SF1, SR1 and SPr) [3]. Amplifications were performed using a StepOne RT-qPCR kit (NEM Luna, cat.no. E3005). Standard curves were generated using plasmids encoding the relevant regions of SARS-CoV2, and the threshold for detection of fluorescence above the background was set within the exponential phase of the amplification curves. CT values from each sample were converted into log10 viral RNA copies/mg tissue according to the standard curve.

Western Blot Analysis:
We used RIPA buffer (ThermoFisher Scientific, cat.no. 89900), supplemented with orthovanadate (New England Biolabs, cat.no. P0758L) and protease inhibitor cocktail (Sigma-Aldrich, cat.no. P8340). Protein concentrations were determined using Pierce 660nm Protein Assay Reagent (ThermoFisher Scientific, cat.no. 22660). Lysates were diluted in 4x Laemmle buffer (Bio-Rad, cat.no. 1610747) and β-mercaptoethanol (ThermoFisher Scientific, cat.no. 21985023), then boiled for 5 minutes at 95°C. Samples were subjected to gel electrophoresis in 4-20% Criterion TGX Stain-Free Protein gels (BioRad, cat.no. 5678095), before being transferred onto PVDF membranes using BioRad Transblot Turbr semi-dry transfer system. Gels were then blocked using Blotting grade blocker (Bio-Rad, cat.no. 170-6404). The following antibodies were used in this study: SARS-CoV2 nucleocapsid (Abcam, cat.no. ab271180), β-actin (abcam, cat.no. ab8224). FABP4 was detected using an in-house antibody produced for the Hotamisligil laboratory by the Dana Farber Antibody Core (clone 351.4.2E12.H1.F12 for capture and HRP-tagged clone 351.4.5E1.H3 for detection). This FABP4 antibody was validated using with protein lysates from FABP4-knockout mice as negative controls and recombinant FABP4 as positive control.

ELISA:
Supernatant of infected cells were mixed with VXL buffer (Qiagen, cat.no. 1069974) in a 1:1 ratio, which has been shown to be sufficient to inactivate SARS-CoV2 [6]. FABP4 secretion was measured by in-house FABP4 ELISA using anti-FABP4 antibodies produced for the Hotamisligil laboratory by the Dana Farber Antibody Core (clone 351.4.2E12.H1.F12 for capture, and HRP-tagged clone 351.4.5E1.H3 for detection) and recombinant human FABP4 (R&D Systems, cat.no. DY3150-05) as a standard. IL-6 was measured following manufacturer's instructions using the human IL-6 Quantikine ELISA kit (R&D Systems, cat.no. D6050).

Real Time Cellular Electronic Sensing (RTCES) Assay:
96-well E-Plates (Acea) were blanked with high glucose DMEM (Gibco) supplemented with 10% FBS (Gibco), 4mM L-Glutamine (ThermoFisher), and 10 U/mL Penicillin Streptomycin (Gibco) to create a baseline reading before seeding the cells at a density of $2.6 \times 10^4$ cells per well. Plates were incubated at room temperature in a biosafety cabinet for 1 hour before being placed into the RTCES Multiplate System (Acea) for 24 hours within a 37°C incubator. Cell impedance was recorded every 15 minutes for every active well. Media was removed and virus

or mock in 2% heat-inactivated FBS, 4mM L-Glutamine, and 10 U/mL Penicillin Streptomycin was added to the cells at a multiplicity of infection (MOI) of 5, 0.5, and 0.05. Plates were returned to the RTCES Multiplate System and incubated at 35°C. RTCES measurements were taken every 15 minutes for 4 days. The median cell index values were determined by fitting the normalized cell index values to a sigmoidal curve using GraphPad Prism software 9.3.1.

Confocal imaging:

Cells were grown and differentiated in collagen-coated glass bottom 35mm imaging dishes, with a no.1.5 coverslip (MatTek, cat.no. P35GCOL-1.5-14-C). At the indicated time points cells were fixed for 20 minutes using 4% paraformaldehyde (PFA; Electron Microscopy Sciences, cat.no. 15710-S) in equal volumes of PBS and growth media. After fixation cells were kept in PBS at 4°C until staining. Prior to staining cells were permeabilized with 0.2% TritonX100 (Sigma-Aldrich, cat.no. 9002-93-1) in PBS for 20 minutes on a slow shaker at room temperature. Cells were then washed twice with PBS. Primary antibodies were diluted 1:200 in PBS and added to the cells overnight at 4°C. After removing the antibodies, cells were washed two to three times with PBS then incubated with the secondary antibody (diluted 1:1000 in PBS) for 1 hours at room temperature. Cells were washed again prior to lipid droplet chemical staining with Bodipy™ 493/503 (ThermoFisher Scientific, cat.no. D3922) diluted 1:1000 in PBS. After another wash, the cells were mounted in mounting media containing Dapi (Vector Laboratories, cat.no. H-1800-10) was added to the cells. The following antibodies were used for this study: anti-SARS-CoV2 nucleocapsid mouse antibody (Cell Signaling Technology, cat.no. 33717), anti-double stranded RNA J2 mouse antibody (Exalpha, cat.no.10010500), anti-calnexin rabbit antibody (Cell Signaling Technology, cat.no. 2679), anti-FABP4 rabbit monoclonal antibody (Abcam, cat.no. ab216708), anti-FABP4 goat polyclonal antibody (Novus, cat.no. AF1443), anti-mouse secondary antibody (Cell signaling technology, cat.no. 4410), anti-rabbit secondary antibody (ThermoFisher Scientific, cat.no. A-11037).

Image analysis was done using ImageJ. To analyze the signal per cell, cells were manually selected and added to the ROI manager by tracing the visible borders of the stained components. After splitting the channels, the background was subtracted, and the intensity histogram of each ROI was analyzed. A set minimum threshold was then defined for each measured signal, and the % area-limited to threshold for each cell was measured. Co-localization analysis was performed using the ImageJ plug-in JACop [7].

Transmission Electron Microscopy (TEM) imaging:

HBE cells were suspended using 0.25% trypsin in EDTA, then fixed with 4% PFA in equal volumes PBS and growth media. Cells were incubated for 20 minutes at room temperature, then centrifuged at 6000 rpm at 4°C for 10min. The pellet was then resuspended in a 1:1 mix of growth media and an EM fixative buffer and incubated at 4°C until TEM processing. The EM fixative buffer contained: [1.25% formaldehyde, 2.5% glutaraldehyde, 0.03% picric acid, 0.05M cocadylate buffer]. For 3D reconstructed airway epithelium organoids, the 4% PFA and fixative buffer were added directly to the apical and basal layers. Samples were washed several times in 0.1M cocadylate buffer, then post fixed with 1% osmiumtetroxide ($OsO_4$)/1.5% potassium ferrocyanide ($K_2FeCN_6$) for 3 hours followed by several washings in water. 1% uranyl acetate in maleate buffer was added for 1 hour, then washed several times in maleate buffer (pH5.2). The samples were then dehydrated in graded cold ethanol series up to 100%, (50%, 70%, 90% then 100%, 10 minutes each) followed by 1 hour incubation in propylene oxide. After the 70%

ethanol dehydration step, 3D airway epithelium cultures were cut out of the insert with a scalpel. Samples were then incubated overnight at 4°C in a 1:1 mixture of propylene oxide and TAAB 812 Resin (TAAB Laboratories Equipment). The following day, samples were embedded in TAAB resin and polymerized at 65°C for 48 hours. 80nm sections were cut with a Leica Ultracut S microtome and placed on formyar-carbon coated slot Cu grids, then stained with 0.2% Lead Citrate and viewed using a JEOL 1200EX electron microscope at 80 kV.
Image analysis was performed using ImageJ software on images acquired at a set magnification and the scale bar was used to determine the pixel/micron ratio. The DMVs were manually selected and added to the ROI manager and the DMV area and X, Y centroid coordinates were measured. To determine the DMV distribution, the distances between the DMV centroids were measured based on their X-Y coordinates and the (mean, median, minimum, and maximum) distances per image were calculated.

Histological imaging:
Human lung biopsies were provided by Koc University Department of Pathology and Yedikule Hospital, Department of Pathology, Istanbul Turkey. After Paraffin embedding and sectioning the samples were deparaffinized under 60°C for 30 minutes then underwent a series of washes with PBS with 0.05% Tween 20. The slides were then incubated for 10 minutes in hydrogen peroxide. Antigen retrieval was performed by manually boiling with 1X citrate (pH 6.0) for 10 minutes. Protein blocking was done for 10 minutes. The cells were then incubated at room temperature with anti-FABP4 antibody (Sigma, cat. no. HPA002188) at a 1:400 dilution in PBS. Secondary antibody was done using anti-rabbit HRP-conjugated antibody (Abcam, cat.no. ab64264). Sections were then counterstained with hematoxylin.
For hamster histology analysis, the left lobe of the lung from each hamster was fixed in 4% PFA for 48 hours prior to transfer out of the BSL3 facility. H/E staining and IHC were done by the Histowiz automated histology platform (https://home.histowiz.com/). The following antibodies were used for the IHC staining anti-FABP4 antibody (Abcam, cat.no. 13979), anti-SARS-CoV2 nucleocapsid antibody (GeneTex, cat.no. GTX635686). Image quantification was performed using QuPath (0.5.0).

Statistical analysis:
For the analysis of human cohort 1, the COVID-19 patient data was combined with the healthy controls, making a total of 326 individuals and 1040 observations, with an average cluster size of 3.2. The main association analyses were performed using the generalized linear mixed model (LMM), regressing the FABP4 concentration on the COVID-19 severity (the reference group consists of the healthy individuals), while accounting for the time of collection post symptom onset, as well as patient-level covariates: age, sex and BMI. Because the estimates from LMM can be biased under model mis-specification, an alternative modeling framework for longitudinal analyses was applied - the Generalized Estimation Equations (GEE) - using the exchangeable correlation structure. Since GEE is robust to incorrectly specified correlation structures within the clusters, we performed sensitivity analyses by comparing the estimates from GEE to those based on the LMM. We observed similar results, which lends support to the usage of LMM. (i.e., indicating the absence of model mis-specification). As a secondary analysis, we replaced COVID-19 severity with oxygen support measures, as a categorical variable, and investigated the association between FABP4 and oxygen support, conditional on the time of collection post symptom onset, age, sex and obesity as covariates. We used the same LMM

model as previously described. We combined the individuals who required nasal canula and mask into one group and treated those with no need for oxygen support as the reference group.

Other statistical analyses were performed using GraphPad Prism version 10.1.1. for MacOS. GraphPad Software, San Diego, California USA, www.graphpad.com. All data are presented as mean ± s.e.m.

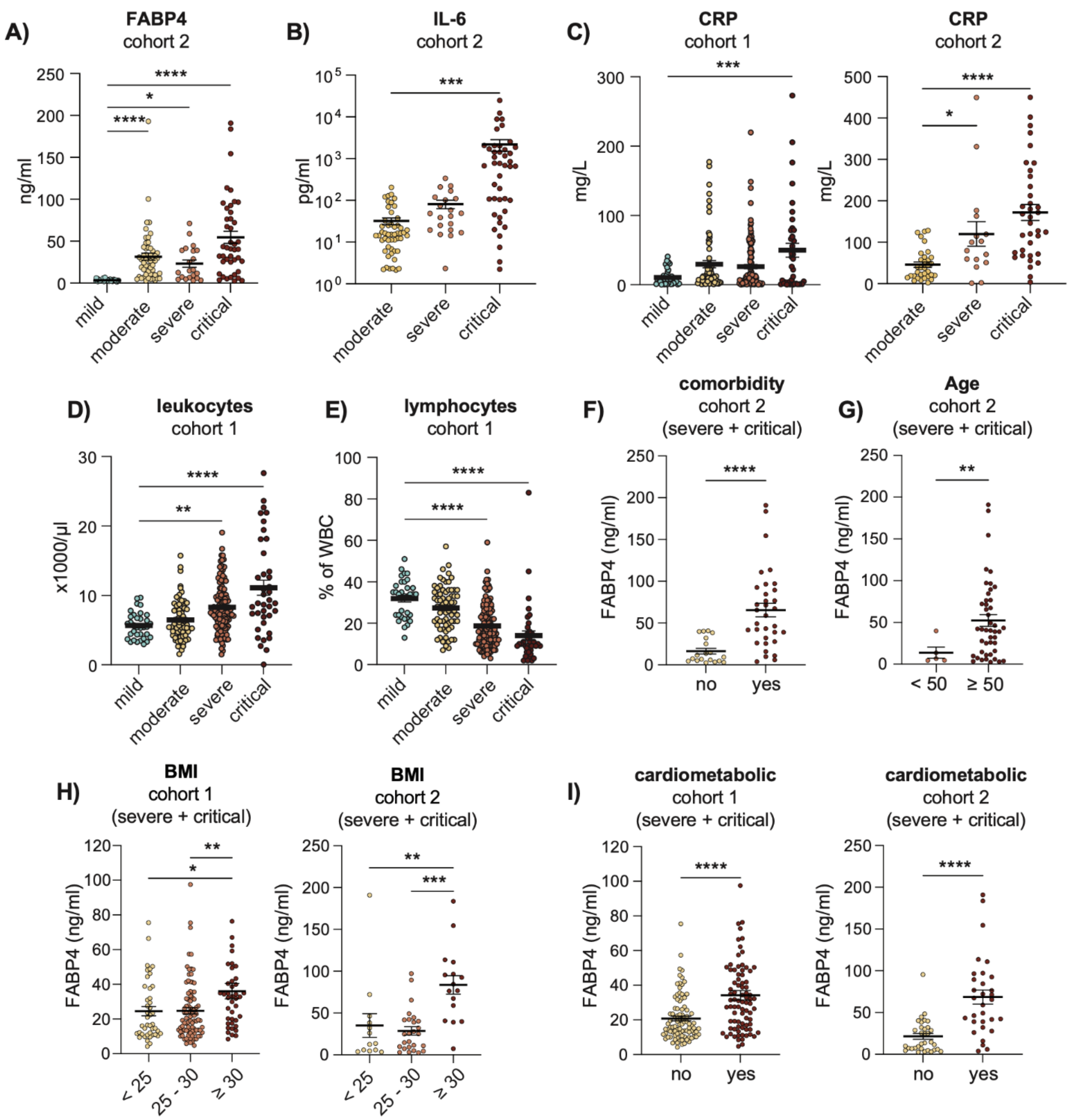


**Supplementary Figure S1: Increase in FABP4 along with biomarkers of COVID-19 disease severity.**

A and B) Maximum concentrations of circulating A) FABP4 and B) IL-6 of cohort 2 of COVID-19 patients stratified based on disease severity. C) Circulating levels of C-reactive protein D) leukocytes and E) lymphocytes of COVID-19 patients measured on the day in which the maximum FABP4 concentration was measured (day post symptom onset). Statistical analysis was performed using one-way ANOVA. F to I) Maximum FABP4 concentration pooled from severe and critically ill patients, stratified based on F) the presence or absence of comorbidities (listed in Tables 3 and S3), G) age, H) BMI and I) the presence or absence of cardiometabolic conditions (diabetes, hypertension, or coronary artery disease). Statistical analysis for F, G and I were performed using Welch's t-test and one-way ANOVA for H. Data are derived from the indicated COVID-19 patient cohort and shown as the mean ± s.e.m.

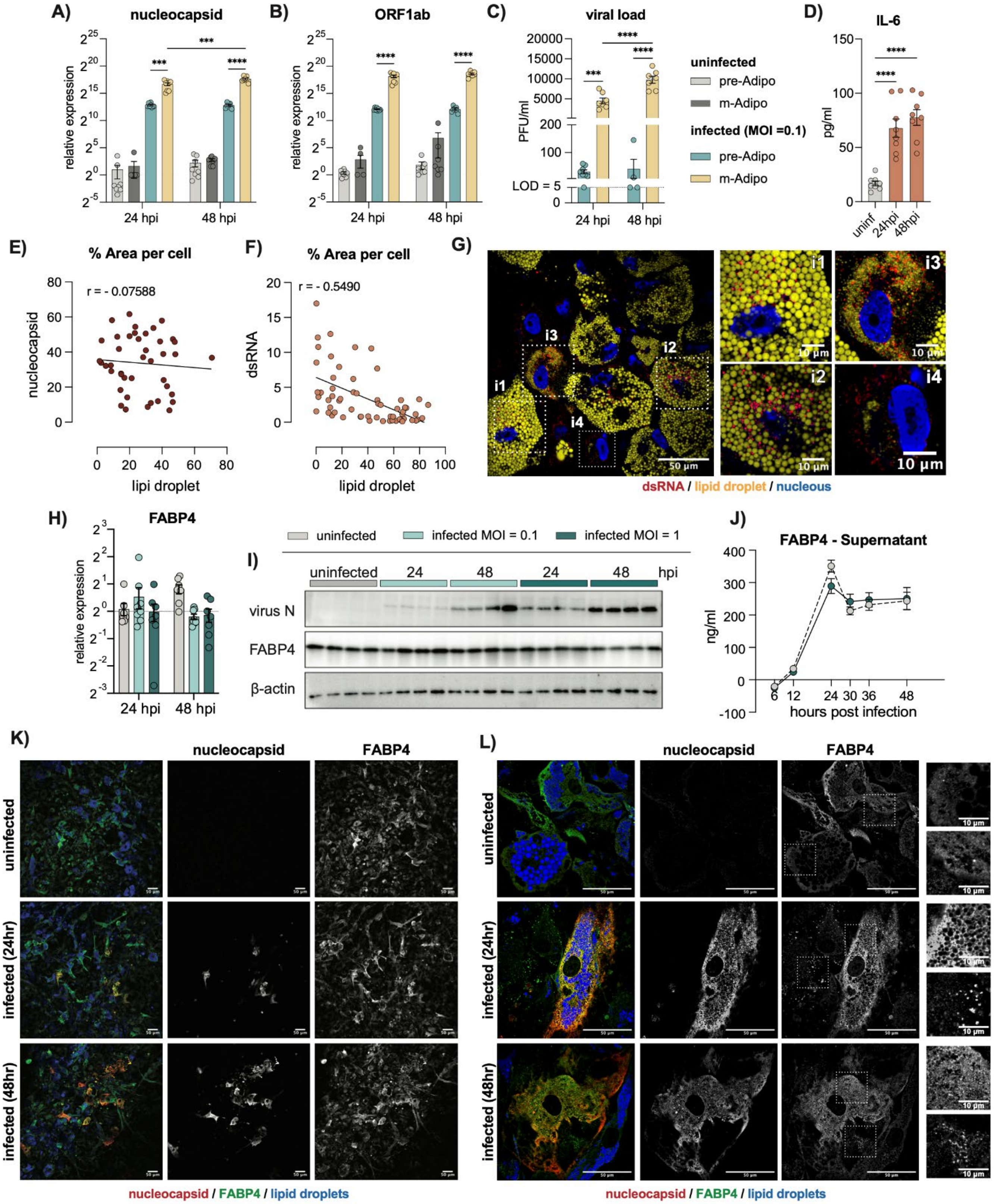
A) nucleocapsid
B) ORF1ab
C) viral load
D) IL-6
relative expression
24 hpi
48 hpi
PFU/ml
LOD = 5
uninfected
pre-Adipo
m-Adipo
infected (MOI =0.1)
pg/ml
uninf
24hpi
48hpi
E) % Area per cell
r = - 0.07588
nucleocapsid
lipi droplet
F) % Area per cell
r = - 0.5490
dsRNA
lipid droplet
G)
dsRNA / lipid droplet / nucleous
H) FABP4
I)
uninfected
infected MOI = 0.1
infected MOI = 1
hpi
virus N
FABP4
β-actin
J) FABP4 - Supernatant
ng/ml
hours post infection
K)
nucleocapsid
FABP4
uninfected
infected (24hr)
infected (48hr)
nucleocapsid / FABP4 / lipid droplets
L)
nucleocapsid
FABP4
uninfected
infected (24hr)
infected (48hr)
nucleocapsid / FABP4 / lipid droplets

**Supplementary Figure S2: FABP4 regulation during SARS-CoV2 infection.**

A to D) pre-adipocytes and mature adipocytes infected with SARS-CoV2 (WA1/2020, MOI: 0.1). A) and B) relative expression of the virus A) genomic (nucleocapsid) and B) sub-genomic RNA (ORF1ab) normalized to β-actin. C) Viral loads measured from supernatant using plaque assay. Data is pooled from two independent experiments, (n=8). Statistical analysis was performed using two-way ANOVA. D) ELISA measuring IL-6 secretion from the supernatant of mature adipocytes in the presence or absence of virus infection (MOI: 0.1). Data are pooled from two independent experiments (n=8). Statistical analysis was performed using one-way ANOVA. E and F) Percent area per cell of lipid droplets relative to E) nucleocapsid or F) dsRNA signal. Pearson correlation is shown as r. G) Representative confocal images of infected mature adipocytes (MOI: 1), strained with dsRNA (red), lipid droplets (yellow), and nucleus (blue, DAPI). Scale bar (50μm) and (10μm) for the magnified areas of interest. (n=3) biological replicates. H) FABP4 gene expression relative to β-actin. Data are pooled from two independent experiments, (n=8). I) Western blots of SARS-CoV2 nucleocapsid and FABP4 protein levels measured from cell lysates of mature adipocytes infected with SARS-CoV2 (MOI: 0.1 or 1). Data are representative of two independent experiments (n=4). J) FABP4 levels, measured by ELISA from the supernatant of infected hTERT cells (MOI: 1). Data are pooled from two independent experiments, (n=12). K and L) Representative confocal images of mature adipocytes stained with nucleocapsid (red), FABP4 (green) and lipid droplets (blue). Scale bar (50μm) and (10μm) for the magnified areas of interest. (n=3) Biological replicates. Data are shown as the mean ± s.e.m.

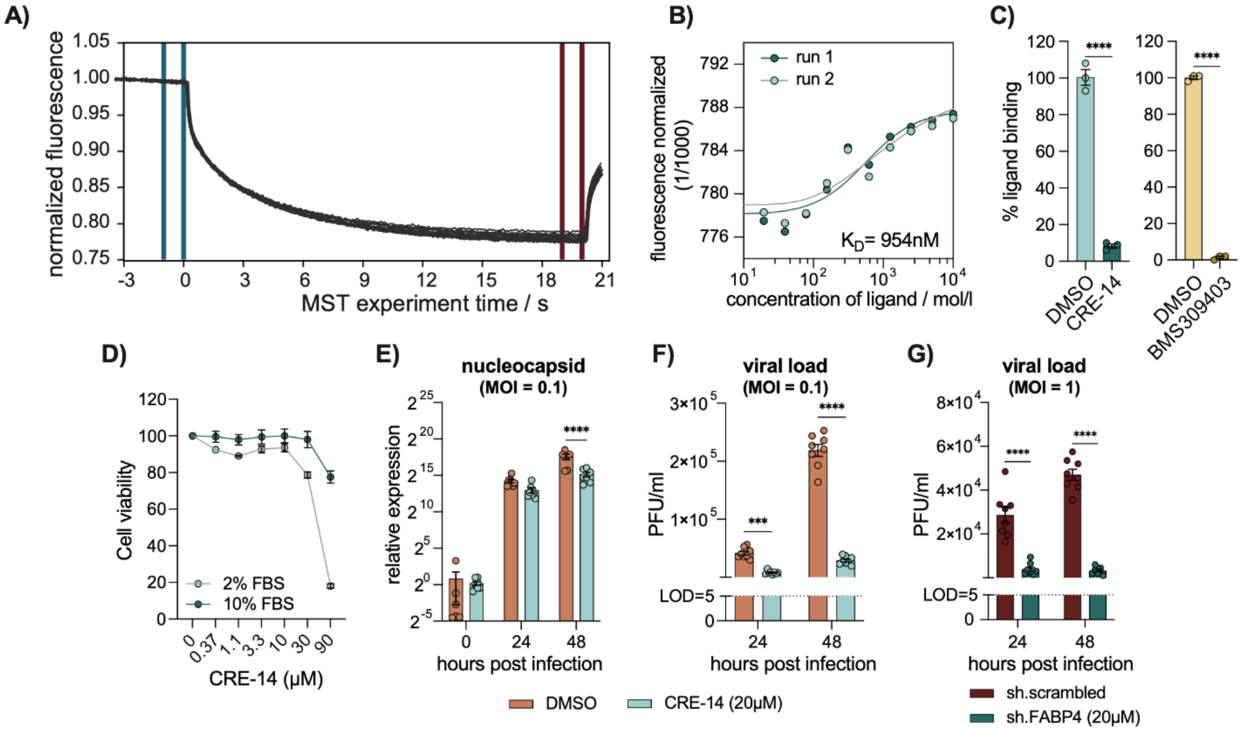


**Supplementary Figure S3: FABP4 deficiency reduces virus titers and cell death following coronavirus infection.**

A) Representative MST time traces with the blue and red regions indicating $F_{cold}$ and $F_{hot}$ respectively, from which the fluorescence measurements were normalized. B) Dose-response curve, showing FABP4 binding to increased concentrations of CRE-14, represented normalized fluorescence. $K_D$ value (954nM) represents the average across two technical runs. C) Percentage of FABP4 bound with fatty acid (BODIPY FL C12) in the presence or absence of CRE-14 or BMS309403. (n=3). D) MRC5 cell viability following administration of titrated doses of CRE-14 at the indicated concentrations of FBS. E and F) Human hTERT adipocytes were infected with SARS-CoV2 (WA1/2020, MOI: 0.1) then treated with either (20µM) CRE-14 or DMSO. E) relative RNA expression of nucleocapsid normalized to β-actin. F) Viral load measured from the supernatant using plaque assay. G) *FABP4*-shRNA knockdown and scrambled controls were infected with SARS-CoV2 (WA1/2020, MOI: 1), and viral titers measured by plaque assay from the supernatants. Data are pooled from two independent experiments, (n=8). Statistical analysis was performed using two-way ANOVA. Data are shown as the mean ± s.e.m.

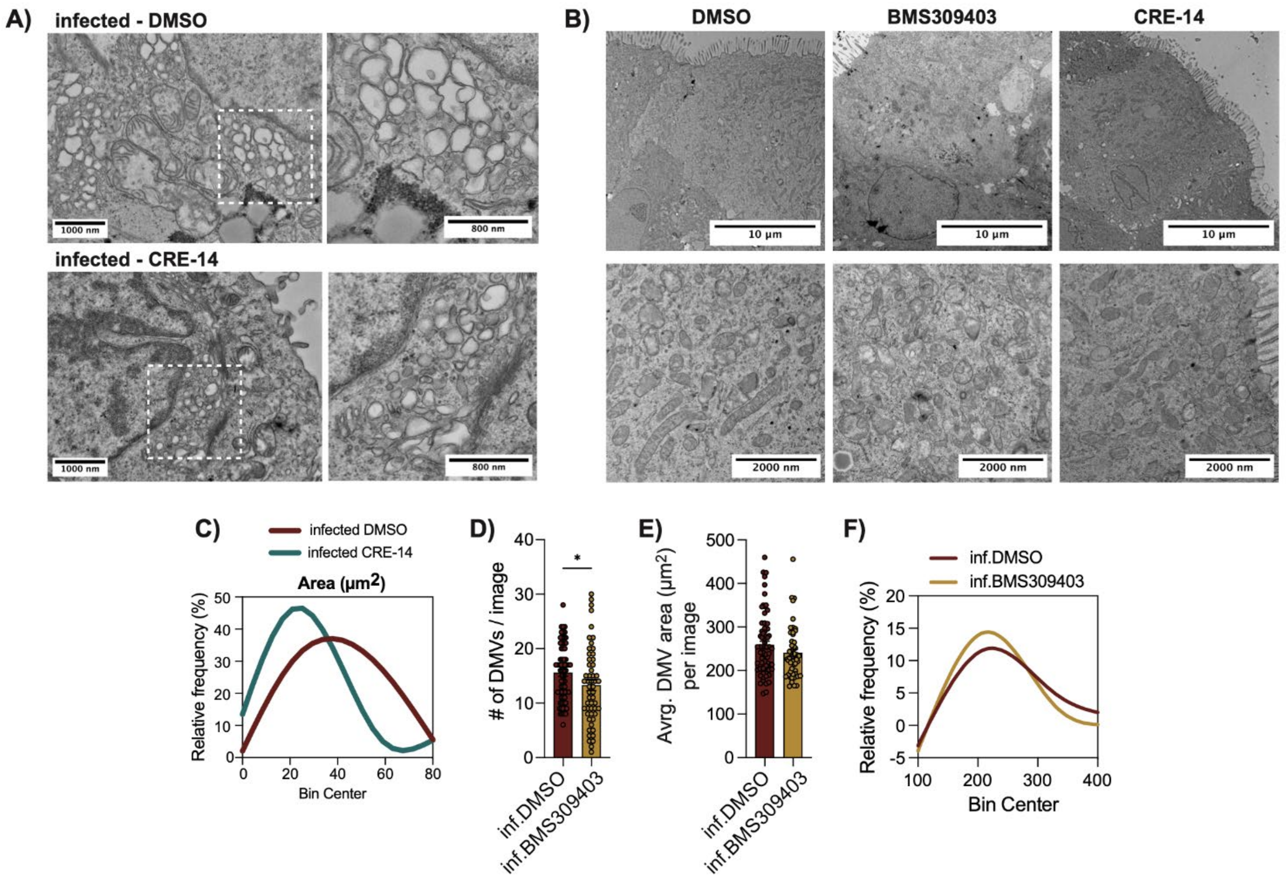


**Supplementary Figure S4: FABP4 inhibition reduces viral titers across various SARS-CoV2 variants.**
A and B) Representative transmission electron microscopy (TEM) images of A) HBE cells 48 hours after infection with SARS-CoV2 (WA1/2020, MOI: 1) and treatment with DMSO or CRE-14 (10µM). (n=3). B) Uninfected reconstructed airway epithelium 3D culture treated with DMSO, BMS309403 or CRE-14. C) Area of double membrane vesicles in infected HBE cells, determined from TEM images in A. Data displayed as the Fit Spline of the percent frequency distribution. D) Number of DMVs per image, E) average DMV area per image and F) its frequency distribution quantified from TEM images reconstructed airway epithelium cultures infected with SARS-CoV2 and treated with DMSO or BMS309403. Statistical analysis is performed using standard t-test. (n=3). Data are shown as the mean ± s.e.m.

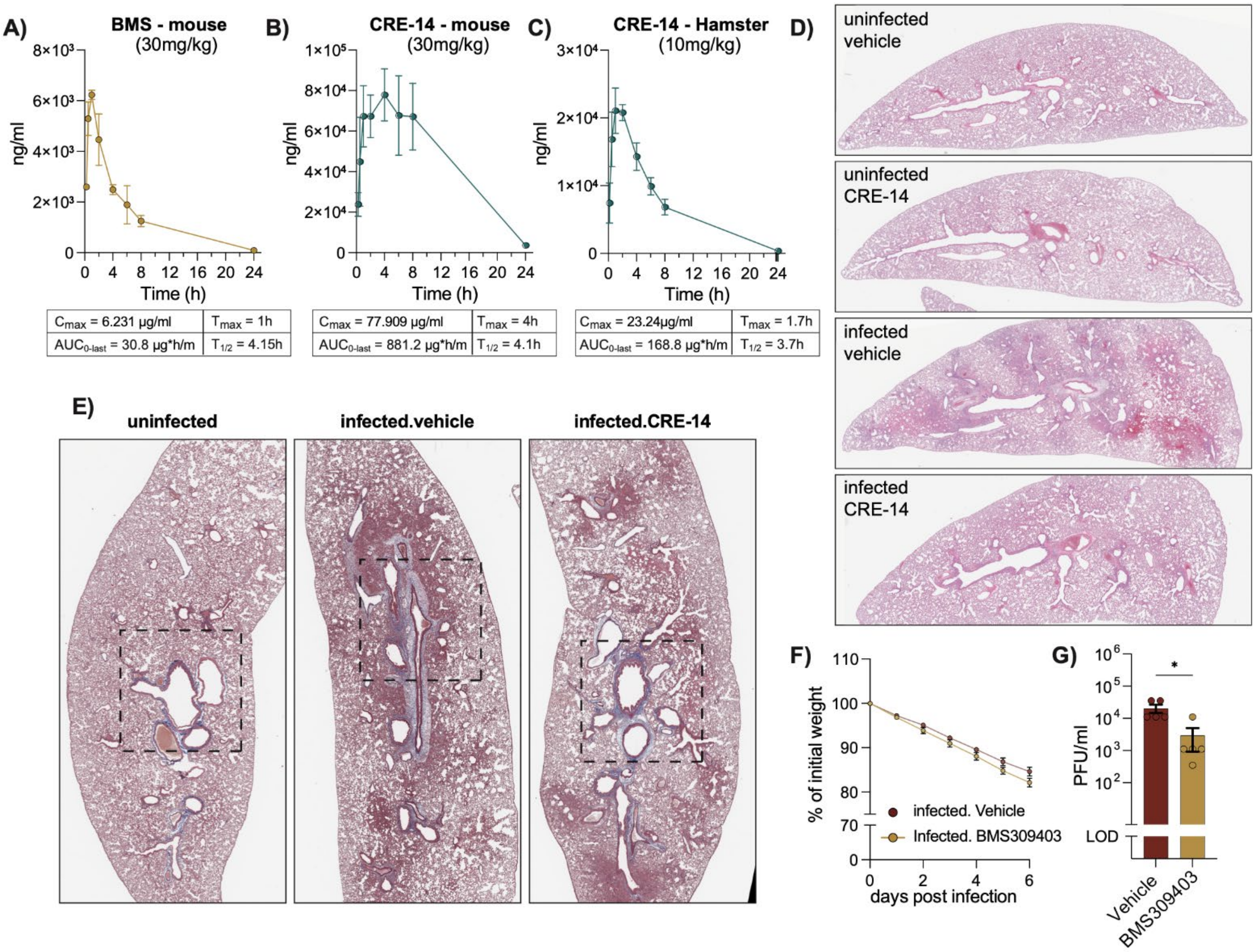


**Supplementary Figure S5: Pharmacokinetics of FABP4 inhibition in mice and hamsters.**
A and B) Circulating concentrations of the BMS309403 and CRE-14 following a 30mg/kg subcutaneous injection in C56BL/6J mice. C) Circulating concentrations of CRE-14 inhibitor in Syrian hamsters following a subcutaneous injection of 10mg/kg. Tables show the ($C_{max}$) maximal concentration, ($T_{max}$) time to reach maximal concentration, ($T_{1/2}$) half-life and ($AUC_{0\text{-last}}$) area under the curve from time zero to the last quantifiable time point. (n=3). D and E) Representative of whole lungs images stained with D) H&E or E) Masson's trichrome across infected and uninfected hamsters with or without CRE-14 treatment (15mg/kg). F) Lung viral titers and G) Body weight loss of Syrian hamsters infected with SARS-CoV2 (Ank1 strain, 100TCID$_{50}$) with or without treatment with 30mg/kg of BMS309403 inhibitor. (n=5). G) Statistical analysis was performed using E) standard t-test. Data are shown as the mean ± s.e.m.
